# The evolution of cooperation in the unidirectional division of labor on a tree network

**DOI:** 10.1101/2023.06.23.546218

**Authors:** Md Sams Afif Nirjhor, Mayuko Nakamaru

## Abstract

Division of labour on complex networks is rarely investigated using evolutionary game theory. We investigate a division of labour where divided roles are assigned to groups on the nodes of a general unidirectional finite tree graph network. From the network’s original node, a task flows and is divided along the branches. A player is randomly selected in each group of cooperators and defectors, who receives a benefit from a cooperator in the upstream group and a part of the task. A cooperator completes their part by paying a cost and then passing it downstream until the entire task is completed. Defectors do not do anything and the division of labour stops, causing all groups to suffer losses due to the incomplete task. We develop a novel method to analyse the local stability in this general tree. We discover that not the benefits but the costs of the cooperation influence the evolution of cooperation, and defections in groups that are directly related to that group’s task cause damage to players in that group. We introduce two sanction systems one of which induces the evolution of cooperation more than the system without sanctions and promote the coexistence of cooperator and defector groups.

## 1 Introduction

According to historical and anthropological material that has been provided in earlier studies regarding our modern human civilization, division of labour was evident in many pre-industrial societies and was connected to their evolution (e.g. Nolan and Lenski, 2011; Durkheim, 1893). The division of labour existed even in civilizations without organized systems like governments (Chauvin and Ozak, 2017). The division of labour results in specialization, which is the foundation of the contemporary, industry-centered civilization. Trade and specialization occur both inside and between industries in the global production networks. Thus, there is a division of labor that is both national and global. As the countries become more and more like each other, more and more intraindustry as well as interindustry specialization is about to occur (Krugman, 1981). The division of labor is a prerequisite for some social systems. One of the examples is supply chain in the economic system which originates from intraproduct specialization (e.g. Krugman, 1981; Fengru and Guitang, 2019). The supply chain have several stakeholders and consists of several suppliers, manufacturers, distributors, retailers, and customers on the basis (Fengru and Guitang, 2019). The multi-layer chain subcontracting system, where a client places the work order with a main contractor, which also places the work order with subcontractors, and this continues until the work reaches the bottom-layer contractors, also depends on intraproduct specialization, thus it is also a case of division of labour (e.g. Tam et al, 2011). The bureaucratic system has a hierarchical structure, based on the division of labor, where the specialization in the specific tasks or official duties leads to the success of the bureaucratic system or organization (e.g. Eisenstadt, 1958; Hinings et. al, 1967; Bolin and Härenstam 2008).

To complete the division of labor, mutual trust and cooperation between groups as well as within groups in the social system is crucial. Even though mutual cooperation is the foundation of human civilization (Turchin, 2016) and our society have prospered through mutual cooperation (Kollock, 1998), individual irrationality, on the contrary, often leads to collective irrationality, and this causes the social dilemma or free riding of defectors. The theoretical and experimental studies on the evolution of cooperation have investigated what mechanisms promote the evolution of cooperation and hinder defectors and free-riders on the basis of the evolutionary game theory.

Nowak (2006) and Rand and Nowak (2013) describe that there are five mechanisms to encourage cooperation; these are kin selection (Hamilton, 1964), direct reciprocity (e.g. Axelrod and Hamilton, 1981; Nowak and Sigmund, 1993, Press and Dyson, 2012), indirect reciprocity (e.g. Sugden, 1986; Nowak and Sigmund 1998, 2005; Nakamaru and Kawata, 2006; Ohtsuki and Iwasa 2004), network reciprocity (e.g. Nowak and May, 1992; Nakamaru et al., 1997, 1998; Ohtsuki et al., 2006; Pacheco et al., 2008; Santos et al., 2008; Su et al., 2022), and group selection (e.g. Sober and Wilson, 1999; Traulsen and Nowak, 2006). In addition to these, the utilization of sanctions to enforce cooperation has also been investigated (e.g. Axelrod, 1986; Sigmund et al., 2001; Boyd et al., 2003; Nakamaru and Iwasa, 2005, 2006; Rand et al., 2010; Sigmund et al., 2010; Shimao and Nakamaru, 2013; Chen et al., 2014, 2015; Sasaki et al., 2015).

The division of labour exists in our society and cooperation is a key there, and the division of labour is studied lately in terms of evolutionary game theory. There are some theoretical works dealing with the division of labor (e.g. Kuhn and Stiner, 2006; Henrich and Boyd, 2008; Nakahashi and Feldman, 2014; Powers and Lehmann, 2014; Roithmayr et al., 2015; Zhang et al., 2018; Qin et al., 2020; Zhang et al., 2020). These studies considered the evolution of the division of labor when there are two or three social roles. Basically, many previous studies assumed a non-structural division of labour among an infinite number of players (e.g. Zhang et al., 2018; Yuan and Meng, 2022). The evolutionary dynamics of the division of labour on a cycle network has also been studied using the division of labor game in which two players, who play different roles from each other, get a higher payoff than two players who play the same role, and it is assumed that each player can choose one of two roles and plays the game with their neighbors on the cycle network (Zhang et al., 2020). The division of labor is a premise of supply chain, which has been investigated especially in management science, and no studies about supply chain with network structure have been done by the evolutionary game theory yet (e.g. Min and Zhou, 2002).

In reality, there are often more than two or three social roles on network structures in the division of labor. However, the division of labor with network structure where different roles are represented by different nodes, is largely unexplored in terms of evolutionary game theory. The linear division of labour was explored by Nakamaru et al. (2018) and Nirjhor and Nakamaru (2023) employing evolutionary game theory. Nakamaru et al. (2018) exemplified the Japanese industrial waste treatment system assuming that there are 3 roles, and Nirjhor and Nakamaru (2023) studied the general case in which there are a finite number of roles. Here, each subtask should be finished in order, consecutively. They selected a finite number of subtasks in an attempt to simplify the model and made the assumption that there are players in each group, where one subtask is assigned to the players of each group. A player in the upstream group randomly chooses a player in the immediate downstream group. If the player in the upstream group is a cooperator, the player does his task by paying a cost of cooperation and passes it to the downstream player who receives the benefit. Then if the upstream player is a defector, the player neither does any task nor passes it to the downstream player. If all players who are chosen from all groups are cooperators, the division of labor is completed, otherwise, it fails. Since the payoffs depend on their tasks and are asymmetric for members of various groups, this system may be represented using the replicator equations for asymmetric games. Through the use of replicator equations for asymmetric games, they investigated whether the two present sanction systems, the defector sanction system and the premier sanction system, can promote the evolution of cooperation. In the premier sanction system, the player in the first group is penalized by the supervisor if defection is discovered in the linear chain. If a defector is discovered under the defector sanction system, they are punished by the supervision. Though both sanction systems are capable of enforcing cooperation more than the system without sanction, when it is practically impossible to monitor and identify defectors the first role sanction system, has been proven to be more successful at ensuring the evolution of cooperation than the defector sanction system. Nirjhor and Nakamaru (2023) found out that the premier sanction system is incapable of creating cooperation when the cooperators are rare initially, however, the defector sanction system is capable of creating cooperation. Nirjhor and Nakamaru (2023) have also found out that the benefit given to a player by a cooperator in the former group has no effect on the evolution of cooperation, and showed that the defector sanction system promotes the coexistence of groups full of cooperators and a group full of defectors when the cost of cooperation increases in the downstream groups.

The division of labor on a tree graph is more common than the linear division of labor in our real world. For example, government in most countries have a system of hierarchy. It has a governmental head or premier, under whom there are several departments, each of which has a head of its own. Each department then breaks down into several sub-departments and it goes until the root level. Therefore, a government system can be considered as a finite tree graph network, which has an origin at the premier and a finite number of branching. Each of the nodes represents a government official. Most governmental action can be considered as a division of labour (Bezes and Le Lidec, 2016), which is ordered by a head and then passes through the downstream nodes and gets fulfilled at some terminal node. Therefore, for an order to be carried out cooperation is very important. Often a single order is carried down and executed by a single linear chain of command which is similar to the linear division of labour which was our previous study (Nirjhor and Nakamaru, 2023). However, to see a governmental system as a whole, a tree graph structure is suitable.

The minimum structure of the supply chain is linear, consisting of suppliers, manufacturers, distributors, retailers, and customers (e.g. Min and Zhou, 2002). However, network-focused models can depict a better image of supply chain than linear chain models (Henderson et al., 2002). The supply chain looks more like a tree than a linear pipeline or chain (Lambert and Cooper, 2000; Cooper et. al, 1997). Therefore, a tree network is capable of depicting the linear network of the supply chain, as well as more general cases. When considering a unidirectional tree network as a supply chain, from the perspective of a player in a terminal node, it is a linear network of the supply chain. However, a player situated in some earlier node can divide the goods along the process links (Min and Zhou, 2002) as well as the labour or responsibility required to improve upon those according to the need. In addition, the multilayered subcontract can be depicted by a tree network (Tam et. al, 2011).

In this paper we study the evolution of cooperation in the division of labour on a general tree graph. No previous models capture all the aspects of a general tree graph in the supply chain (e.g. Min and Zhou, 2002), and our study challenges this problem by means of the evolutionary game theory.

## 2 Baseline Model and Results

### 2.1 Model Assumption

We take a model where the whole task is divided and assigned to the groups who are present in the nodes of a connected directed tree graph. The model structure is shown in figure 1. In this model a task is always passed from the upstream to the downstream, never from downstream to upstream, hence, this is unidirectional. There is a unique central node in this graph, and from there branching starts. **G** is the set of nodes in the tree graph (figure 1). Each of the nodes has a group of players, each group consists of cooperators and defectors, and the group population is infinite. *p* is the index of the original node which represents the premier group. In the beginning, the whole task, which can be a service or development of a product, is assigned to the original group, where a player is randomly selected, who gets a benefit *b*_*p*_. The player receives the benefit for receiving the task and it can be considered as the value of the task (Nirjhor and Nakamaru, 2023). For example, in the multi-layered contract development system, the orderer pays the contract money, *b*_*p*_, to the contractor, which pays the money to the subcontractors and it continues when the terminal sub-subcontractors receive the money and do their tasks. In the industrial waste disposal system it can be considered as the benefit from the product, from which the waste was produced (Nakamaru et. al, 2018).

**Figure 1:**
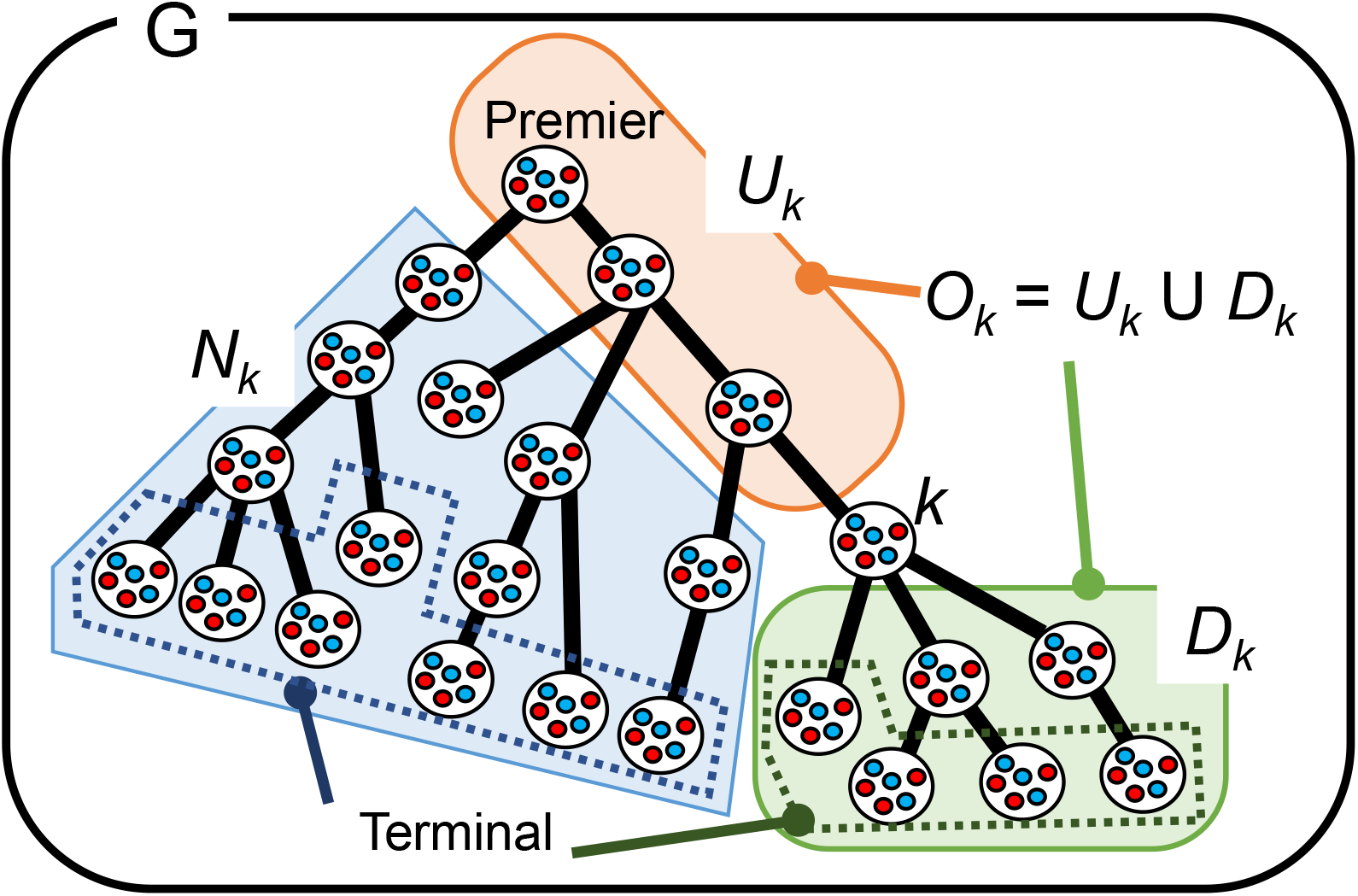
Division of labour in a downstream tree graph. *k* is the focal group. The upstream of *k, U*_*k*_ is a linear network from the premier, the downstream of *k, D*_*k*_ is a tree graph network. *U*_*k*_ and *D*_*k*_ makes the *O*_*k*_. *N*_*k*_ is created with the groups that are not present in *O*_*k*_.

If the chosen player in a group is a cooperator, they pay a cost of cooperation *x*_*p*_ to improve upon the task. Then they divide the task and then pass the task to one or more downstream branches. Then each player who is chosen randomly from each of the receiving downstream groups, and also receives a benefit, as each has received the task. Table 1a shows the payoff of a cooperator in the premier group when all players chosen from all downstream groups are cooperators: *b*_*p*_ − *x*_*p*_ (table 1a).

**Table 1a:** The payoff matrix in Baseline system for Premier

| Cases | All being cooperator except the premier | A defector after the premier |
| --- | --- | --- |
| Premier cooperator | $b_p - x_p$ | $b_p - x_p - g_{op}$ |
| Premier defector | $b_p - g_p$ | $b_p - g_p$ |

**Table 1b:** The payoff matrix in the baseline system for *k* ≠ *p*

| Cases | All being cooperator in $\mathbf{O}_k$ | A defector in $U_k$ | A defector in $D_k$ |
| --- | --- | --- | --- |
| $k$ cooperator | $b_k - x_k - g_{nk}$ | $-g_k - g_{nk}$ | $b_k - x_k - g_{ok} - g_{nk}$ |
| $k$ defector | $b_k - g_k - g_{nk}$ | $-g_k - g_{nk}$ | $b_k - g_k - g_{nk}$ |

**Table 1c:**
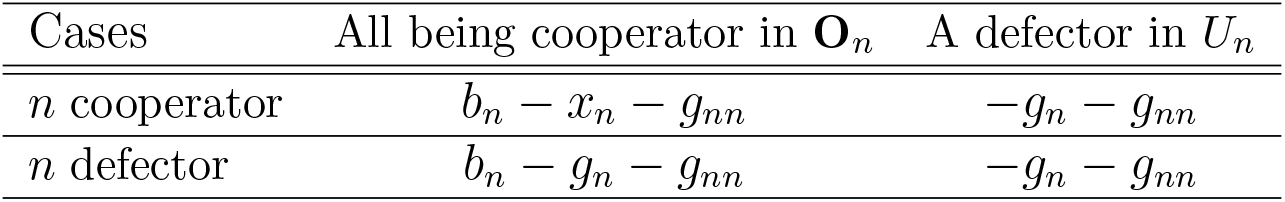
The payoff matrix in the baseline system for *n≠ p* in terminal

| Cases | All being cooperator in $\mathbf{O}_n$ | A defector in $U_n$ |
| --- | --- | --- |
| $n$ cooperator | $b_n - x_n - g_{nn}$ | $-g_n - g_{nn}$ |
| $n$ defector | $b_n - g_n - g_{nn}$ | $-g_n - g_{nn}$ |

If a defector is chosen from the premier group, the defector does not produce any benefit paying a cost of cooperation, and the division of labor does not start. As a result, the player from the premier group will suffer from the loss, *g*_*p*_, which can be interpreted as the damage or the bad reputation from the incomplete tasks of the whole system (table 1a). Then, a player in the downstream group *k* (*k* ≠ *p*) suffers from the loss, *g*_*k*_, and it is assumed that *g*_*p*_ = Σ*g*_*j*_, where *j*s are the groups present in the immediate branching of the premier. We will explain the assumption of the loss caused by defection later.

Even though a cooperator is chosen from the premier group, the cooperator suffers from the loss, *g*_*op*_ if a defector is chosen in the downstream groups in *D*_*p*_ and the division of labor stops there (table 1a). To define *D*_*p*_, we have to define *O*_*k*_, *U*_*k*_, *D*_*k*_, and *N*_*k*_ with respect with group *k* at first (figure 1). *O*_*k*_ is the set of nodes that create *k*’s connection with the premier and are present on the branches originating from *k* if *k* is not premier (figure 1). *U*_*k*_ is the set of the upstream nodes of *O*_*k*_ with respect to *k* and these nodes are in a linear division of labour with respect to *k. D*_*k*_ is the set of the downstream nodes of *O*_*k*_ with respect to *k*. Therefore, *O*_*k*_ = *U*_*k*_∪*D*_*k*_ (figure 1). When a defector is chosen from group *k*, during that particular task, *D*_*k*_’s groups become neutral, as they do not have a choice. *N*_*k*_ = **G** − (*O*_*k*_∪{*k*}) is the set of nodes that do not have the direct or indirect interaction with *k*.

If *k* is premier or *p*, both *U*_*p*_ and *N*_*p*_ do not exist, and *D*_*p*_ is equal to *O*_*p*_. If *k* is terminal or *t, D*_*t*_ does not exist, *U*_*t*_ is equal to *O*_*t*_, and *N*_*t*_ = **G** − (*U*_*t*_∪{*t*}).

We consider *k* is the index of our focus group, and this focus group can be any group in the graph. The value, *b*_*k*_, is the benefit of the player chosen from the group *k*, given by the cooperator in the nearest upstream group *u* with paying a cost of cooperation *x*_*u*_. The benefit can also be considered as the value of the task. If the player in group *k* is also a cooperator, he pays the cost of cooperation *x*_*k*_ to produce a new task or value, *b*_*kk*_, and then passes it to his group’s branch/branches if group *k* is not terminal. The net benefit of the cooperator in group *k* is *b*_*k*_ − *x*_*k*_ if all other players are cooperators. If group *k* has two nearest downstream groups, for example, and they are named group A and group B. A cooperator in the group *k* gives a benefit to each player in two groups. There are two possible assumptions: the cooperator in group *k* will give a benefit *b*_*A*_ to a player in group A and *b*_*B*_ in a player in group B, where, (i) *b*_*kk*_ = *b*_*A*_ + *b*_*B*_, or (ii) *b*_*kk*_ = *b*_*A*_ = *b*_*B*_. If the benefit is a divisible good such as a product or money, it should be divided and (i) can be applied. If the benefit is an indivisible good such as a service, (ii) can be applied. Both of the cases can be covered in this model, as we shall see in the expansion of the model that the benefit itself shall disappear from the dynamics.

This continues until the terminals unless a defector is selected. If a defector is selected in group *k*, the defector just receives the benefit, *b*_*k*_, from the cooperator in the upstream group and does not do a task paying the cost of cooperation, does not pass the task to his downstream. Hence, the division of labour stops there, that particular task is not completed and everyone in every group bears the loss, *g*_*k*_.

Here, we explain our assumption about the losses caused by defectors. We assume that the loss to everyone if a defector is chosen in group *k*, is *g*_*k*_, which is divided into the immediate branching of group *k*; *g*_*k*_ = Σ_*j*_ *g*_*j*_ where *j*s are the immediate branching of *k*. Each player suffers from the same losses caused by defectors in the whole system. However, from the viewpoint of group k, the losses a player in group *k* suffers from are classified into three types; the self-inflicted loss (*γ*_*k*_), the potential loss (*g*_*ok*_), and the loss caused by *N*_*k*_ (*g*_*nk*_). The total loss to a player in group *k* is *γ*_*k*_ + *g*_*ok*_ + *g*_*nk*_.

The self-inflicted loss *γ*_*k*_ is *g*_*k*_ when a defector is chosen in group *k*, and zero when a cooperator is chosen. When defection occurs in *U*_*k*_, *γ*_*k*_ = *g*_*k*_; we assume that when the downstream player suffers from the loss caused by the upstream defector, the loss of the downstream player is the same as what he would have suffered by his defection. This means, when a player in a group in *U*_*k*_ chooses defection, the product or service which would have been produced or done by the player in group *k*, was not produced or done, so the loss born by the player in group *k* is the same as *g*_*k*_.

When the player in group *k* has already cooperated, some part of *D*_*k*_, may cooperate and some part may not. Then, the cooperator in group *k* suffers from the loss by defection in *D*_*k*_. This loss is called the potential loss to cooperators in the upstream groups, *g*_*ok*_, which is defined as the combined loss through branching in *D*_*k*_; *g*_*ok*_ = Σ *δ*_*j*_(*t*)*g*_*j*_ where *j* ∈ *D*_*k*_ and *δ*_*j*_(*t*) is 1 when a chosen player in group *j*s are defectors in *D*_*k*_ at time *t*, and zero otherwise. When the cooperator is in the group *k* and all the chosen players in the groups of *k*’s immediate branching are defectors, then *g*_*ok*_ = Σ *g*_*j*_ = *g*_*k*_. Otherwise, *g*_*ok*_ < *g*_*k*_. This condition makes the choice of branching for a player impartial, as the distributed risk of defection is the same as or lower than defection by oneself.

We also assume that any player in group *k* suffers from the loss caused by *N*_*k*_, *g*_*nk*_. The value *g*_*nk*_ includes the losses caused by defectors in *N*_*k*_. In addition, when there is a defection in a group *l* ∈ *U*_*K*_, that also has branches in *N*_*k*_, the loss *g*_*l*_ damages everyone in every group. A part of this loss comes to group *k* as the self-inflicted loss *g*_*k*_, the rest of it flows in *N*_*k*_ and gives rise to the self-inflicted losses of players in groups those in *D*_*l*_ ∩ *N*_*k*_. To calculate this loss we take the summation of the self-inflicted losses of the terminals of *D*_*l*_ ∩ *N*_*k*_. In other words, *g*_*nk*_ also includes the self-inflicted losses of the groups which are in the terminal of *N*_*k*_ and had a defector in their upstream that intersects with *U*_*k*_. In sum, the mathematical definition: *g*_*nk*_ = Σ *g*_*j*_ + *g*_*m*_, where *j* ∈ *N*_*k*_ and *j* are defectors when cooperators are selected in *U*_*k*_ ∩ *U*_*j*_, and *m* ∈ *N*_*k*_ and *m* are terminals when a defector is selected in *U*_*k*_ ∩ *U*_*m*_.

Figure 2 and table 2 show the example of the losses in the 2-regular 2-branched directed tree graph. Here if the group index is *k*_*j*_, the loss is shown as 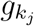. The relationship between losses is 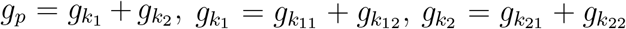. The self-inflicted loss of a cooperator in the premier group is zero. The potential loss that a cooperator in the premier group suffers from defection in the downstream groups, *D*_*p*_, is *g*_*op*_, which is the sum of the losses when defectors are selected after the premier group, 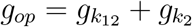 in figure 2. *g*_*np*_ is zero because *N*_*p*_ is empty. In group *k*_11_, the self-inflicted loss is zero, the potential loss is zero, and 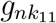 is 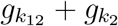 because defection occurs in both groups *k*_12_ and *k*_2_ in 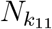. Table 2 shows the three types of losses of other groups in figure 2.

**Table 2:** Losses to the players in the groups in figure 2

| Group $k$ | state | $\gamma_k$ | $g_{ok}$ | $g_{nk}$ | relative equilibrium |
| --- | --- | --- | --- | --- | --- |
| Premier | Cooperation | 0 | $g_{k_{12}} + g_{k_2}$ | 0 | $mix_{D_p}$ |
| $k_1$ | Cooperation | 0 | $g_{k_{12}}$ | $g_{k_2}$ | $mix_{D_{k_1}}$ |
| $k_{11}$ | Cooperation | 0 | 0 | $g_{k_{12}} + g_{k_2}$ | $allc_{k_{11}}$ |
| $k_{12}$ | Defection | $g_{k_{12}}$ | 0 | $g_{k_2}$ | $mix_{U_{k_{12}} \cup \{k_{12}\}}$ |
| $k_2$ | Defection | $g_{k_2}$ | 0 | $g_{k_{12}}$ | $mix_{U_{k_2} \cup \{k_2\}}$ |
| $k_{21}$ | Neutral | $g_{k_{21}}$ | 0 | $g_{k_{12}} + g_{k_{22}}$ | $mix_{U_{k_{21}} \cup \{k_{21}\}}$ |
| $k_{22}$ | Neutral | $g_{k_{22}}$ | 0 | $g_{k_{12}} + g_{k_{21}}$ | $mix_{U_{k_{22}} \cup \{k_{22}\}}$ |

**Figure 2:**
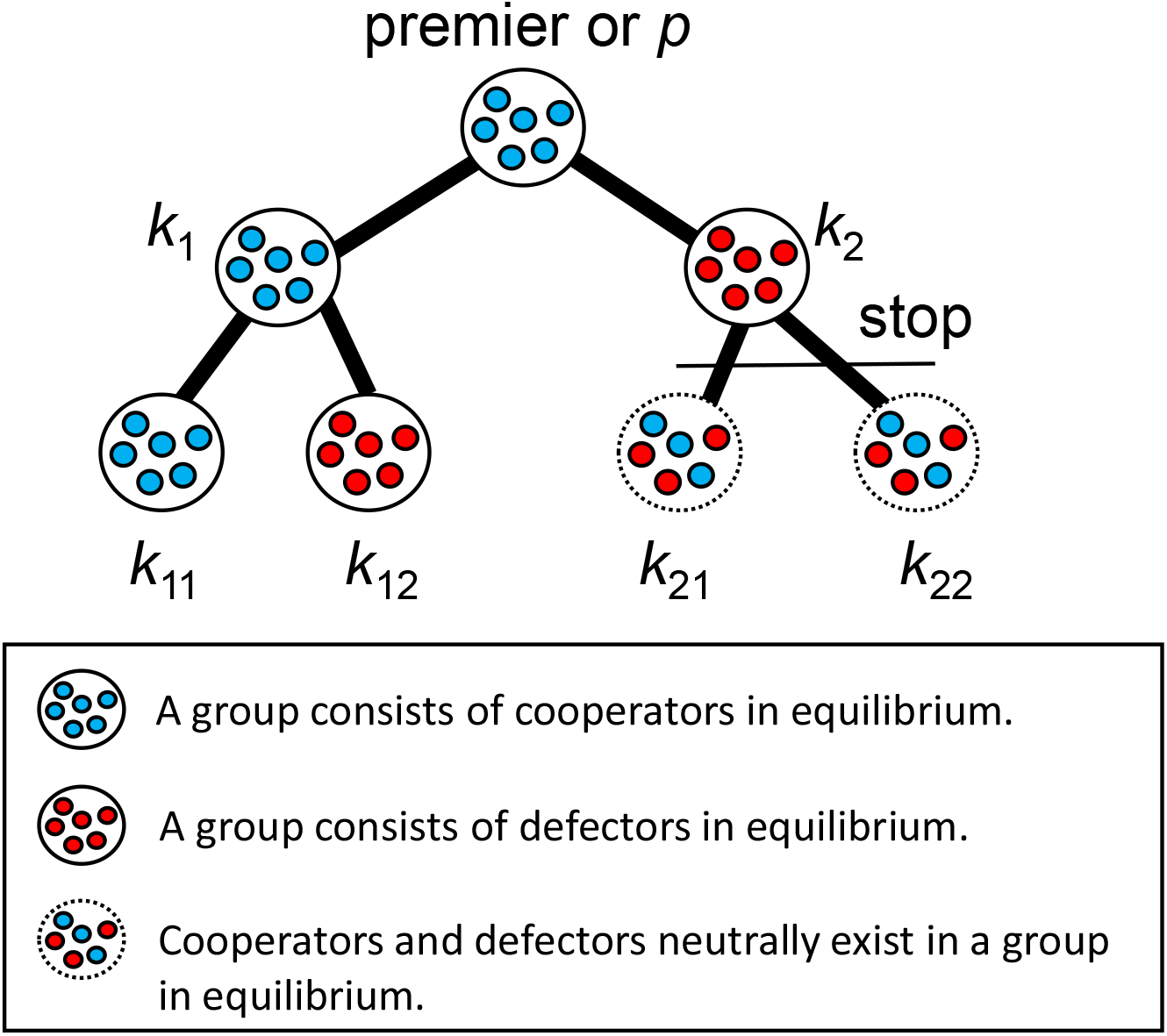
The relative image of the equilibria and the loss distribution in the system of 2-regular 2-branched directed tree graph. The premier group name is *premier* or *p*, and the group name except the premier group is *k*_*i*_ where *i* is 1, 2, 11, 12, 21, or 22. The groups *p, k*_1_, and *k*_11_ are groups consisting of cooperators in equilibrium. The groups *k*_12_ and *k*_2_ are groups consisting of defectors in equilibrium. The groups *k*_21_ and *k*_22_ are groups where cooperators and defectors neutrally exist. Table 1 shows the loss to a player in all groups.

Based on our assumption mentioned above, we can calculate the payoff of either a cooperator or a defector in group *k* (table 1); tables 1a, 1b and 1c are when *k* = *p, k* is neither *p* nor terminal, and *k* = *n* is a terminal, respectively. The parameters are shown in table 3.

**Table 3:** Parameters

|  |  |
| --- | --- |
| $\mathbf{G}$ | Set of nodes in the tree graph |
| $k$ | Index of nodes |
| $p$ | Index of the first or original node, called the premier |
| $O_k$ | Set of nodes which create $k$'s connection with the premier and are present on the branches of $k$ |
| $N_k$ | $\mathbf{G} - (O_k \cup \{k\})$ |
| $U_k$ | Set of the upstream nodes of $O_k$ with respect to $k$ |
| $D_k$ | Set of the downstream nodes of $O_k$ with respect to $k$ |
| $c_{ok}$ | Probability of all the players in all the groups of $\mathbf{O}_k$ |
| $d_{uk}$ | Probability of a defector in $U_k$ |
| $d_{dk}$ | Probability of defector(s) in the downstream of group $k$ in $D_k$ |
| $g_k$ | Loss to everybody if a defector is chosen in group $k$ |
| $g_{ok}$ | the potential loss in $D_k$ |
| $g_{nk}$ | the loss caused by $N_k$ |
| $x_k$ | Cooperation cost of a player in group $k$ |
| $b_k$ | Benefit of a player in group $k$ |
| $k_c$ | The frequency of cooperator in group $k$ |
| $k_d$ | The frequency of defector in group $k$ |
| $f$ | Amount of punishment |
| $\rho$ | Probability of catching a defector, $\rho$ is considerably low |

### 2.2 Replicator equations for asymmetric games

If we assume that each player imitates the behaviour of others with a higher payoff in the same group, we can apply the replicator equations for asymmetric games. To calculate the expected payoff of players in the replicator equations of asymmetric games, three parameters of probability are defined; 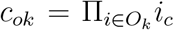 is the probability of all the selected players in all the groups of *O*_*k*_ being cooperators, where *i*_*c*_ is the frequency of cooperators, and *i*_*d*_ is the frequency of defectors in the group *i*. Here, *i*_*c*_ + *i*_*d*_ = 1. 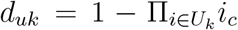 is the probability of a defector being selected in the groups of *U*_*k*_. 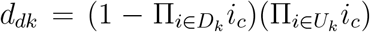 is the probability of defector(s) being selected in the groups which are downstream of *k*, or in other words in *D*_*k*_ there is a defector or defectors. We find, *c*_*ok*_ + *d*_*uk*_ + *d*_*dk*_ = 1. When *k* is the premier, if the player from *k* is a cooperator, and all the players who are selected from other groups are also cooperators, then the payoff of the player in *k* is *b*_*p*_ − *x*_*p*_, as they receive the benefit *b*_*p*_ and pay the cost of cooperation *x*_*p*_, to produce a product or do a task (table 1a). If there are one or more defectors in *k*’s downstream then the payoff is *b*_*p*_ − *x*_*p*_ − *g*_*op*_, as the combined loss due to defection *g*_*op*_ will also be born (table 1a). Therefore when the player from the premier group is a cooperator, then their expected payoff, Π_*cp*_, is *c*_*op*_(*b*_*p*_ − *x*_*p*_) + (1 − *c*_*op*_)(*b*_*p*_ − *x*_*p*_ − *g*_*op*_) (see table 1a). If the player from the premier group is a defector then their expected payoff, Π_*dp*_, is *b*_*p*_ − *g*_*p*_ (see table 1a) as he receives the benefit *b*_*p*_ but does not pay the cost of cooperation. However, due to his defection he needs to bear the loss *g*_*p*_. As he is in the premier group, his defection leads to the linear division of labour being stopped. So, the latter groups’ player’s strategy does not have any effect on his payoff, when he is a defector. Therefore, the replicator equation of a cooperator in the premier group is,

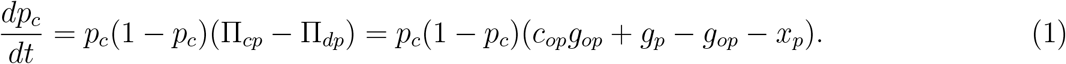

When *k* is neither the premier nor a terminal, if the player from *k* is a cooperator, and all the players who are selected from other groups in *O*_*k*_ are also cooperators, then the payoff of the player in *k* is *b*_*k*_ − *x*_*k*_ − *g*_*nk*_, as they receive the benefit *b*_*k*_, pay the cost of cooperation *x*_*k*_, and also bear the loss *g*_*nk*_ due to the possible defections in *N*_*k*_ (table 1b). If there are one or more defectors in *k*’s downstream then the payoff is *b*_*k*_ − *x*_*k*_ − *g*_*ok*_ − *g*_*nk*_, as the combined loss due to defection *g*_*ok*_ will also be born (table 1b). If there is a defector in the *U*_*k*_, then the task does not reach *k*, so in that case he only bears the loss *g*_*k*_ and their payoff becomes −*g*_*k*_ − *g*_*nk*_. So, the expected payoff of a cooperator in group *k*, Π_*ck*_, is *c*_*ok*_(*b*_*k*_ − *x*_*k*_) − *d*_*uk*_*g*_*k*_ + *d*_*dk*_(*b*_*k*_ − *x*_*k*_ − *g*_*ok*_) − *g*_*nk*_. If the player from *k* is a defector and all the chosen players in *O*_*k*_ are cooperators, or there are defectors to be chosen in *D*_*k*_, then his payoff is *b*_*k*_ − *g*_*k*_ − *g*_*nk*_ as they receive the benefit *b*_*k*_ but do not pay the cost of cooperation. However, due to their defection, they need to bear the loss *g*_*k*_. If there is a defector in the *U*_*k*_, their payoff is the same as being a cooperator, because they do not get a chance to play their strategy. So, when a player in group *k* is a defector, the expected payoff, Π_*dk*_, is *c*_*ok*_(*b*_*k*_ − *g*_*k*_) − *d*_*uk*_*g*_*k*_ + *d*_*dk*_(*b*_*k*_ − *g*_*k*_) − *g*_*nk*_. Therefore, the replicator equation of a cooperator in group *k* is,

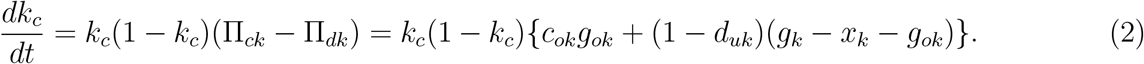

When *n* is a terminal group, the division of labour is effectively a linear network from the point of view of *n*. However, the aspect of possible defections in *N*_*n*_ needs to be considered. If the chosen player from *n* is a cooperator and players chosen from all the other groups in *O*_*k*_ are also cooperators, then their payoff is *b*_*n*_ − *x*_*n*_ − *g*_*nn*_. If there is a defector chosen in any of the groups of *O*_*n*_, then the task does not reach *n*, so their payoff in the terminal group becomes −*g*_*n*_ − *g*_*nn*_, regardless of them being a cooperator or a defector. When all the players chosen from all the other groups in *O*_*n*_ are cooperators, however, the player chosen in *n* is a defector, then his payoff is *b*_*n*_ − *g*_*n*_ − *g*_*nn*_, as he will have to bear the loss, *g*_*n*_ due to his own defection (table 1c).

Hence, a cooperator’s expected payoff from a terminal group *n*, Π_*cn*_ is *c*_*on*_(*b*_*n*_ − *x*_*n*_) − *d*_*un*_*g*_*n*_ − *g*_*nn*_. When the player from the terminal *n* is a defector, his expected payoff, Π_*dn*_ is *c*_*on*_(*b*_*n*_ − *g*_*n*_) − *d*_*un*_*g*_*n*_ − *g*_*nn*_. Therefore, the replicator equation of a cooperator in the terminal group is,

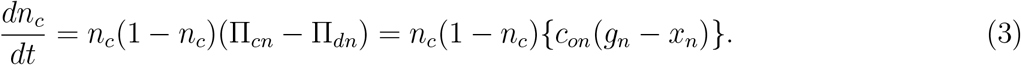

Here, the benefit *b*_*k*_ given by a cooperator of the upstream as well as the term *g*_*nk*_ which represents the loss caused by *N*_*k*_ are both cancelled in equations (1)-(3). Therefore, they do not have any effect on the dynamics. We will show that two values, *b*_*k*_ and *g*_*nk*_, are also cancelled out when the sanction system is introduced in the equations (Section 3 and Appendices B and C).

### 2.3 Results

We do the stability analysis of three sorts of equilibrium for each *O*_*k*_∪{*k*} (figure 3).

**Figure 3:**
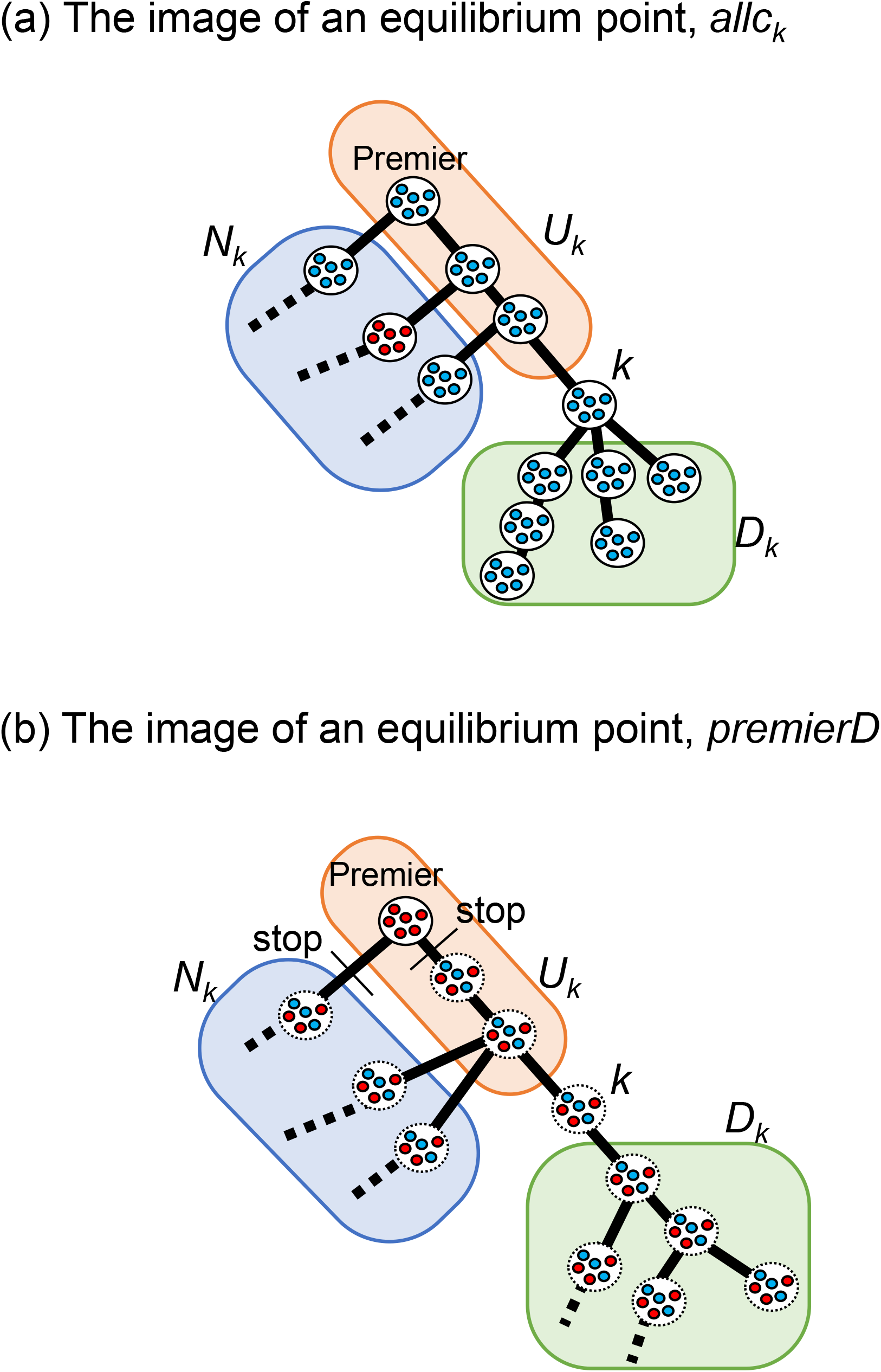
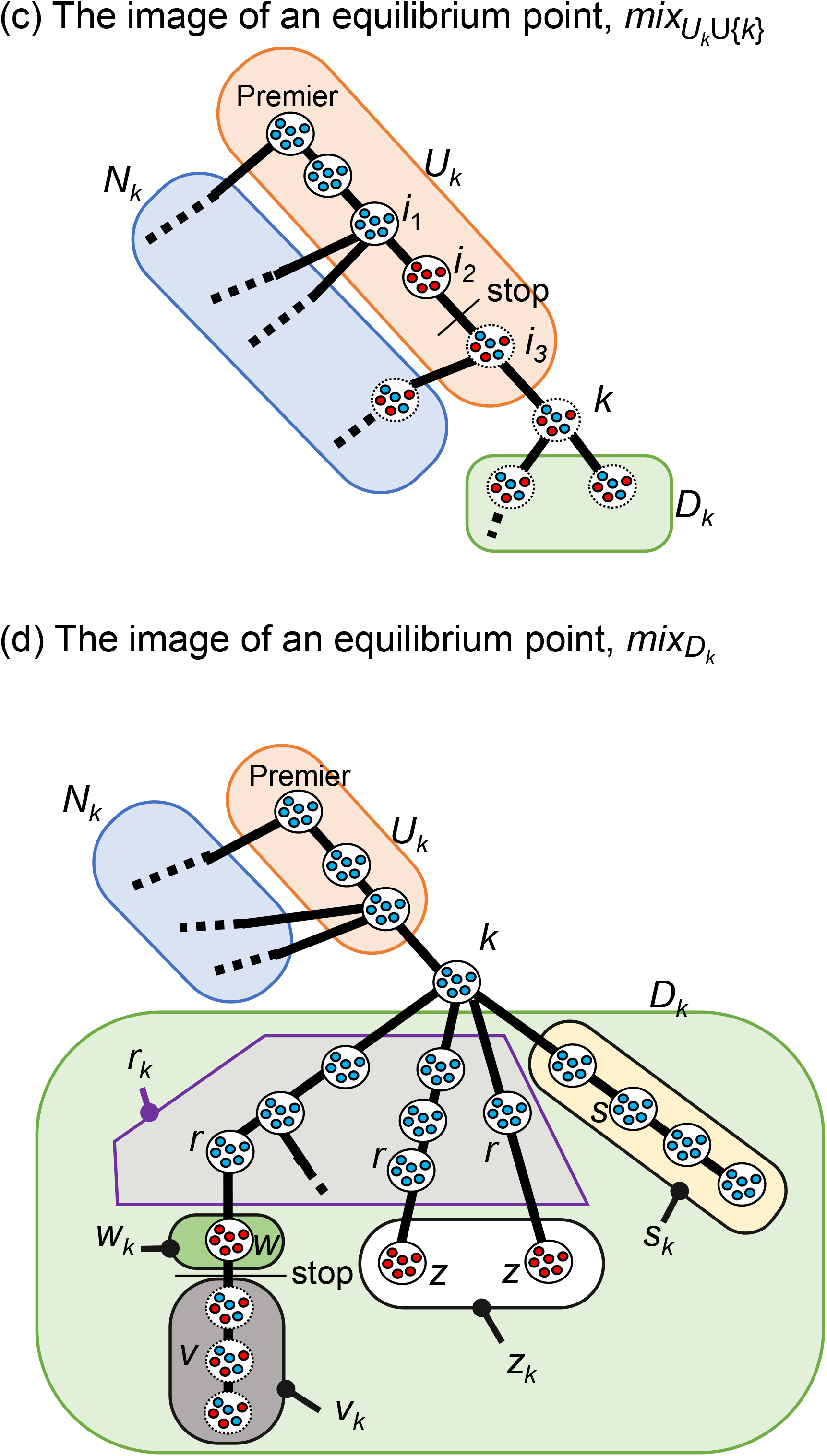
The equilibria in the system from the focal group *k*. (a) is showing the *allc*_*k*_, (b) is showing the *premierD*, (c) is showing the 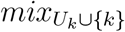, and (d) is showing the 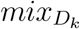.

The all-cooperation equilibrium is defined as that everyone in every group is a cooperator in *O*_*k*_∪{*k*}, which is hereafter called *allc*_*k*_ (figure 3a). The premier group defection equilibrium is defined as that everyone in the premier group *p* is a defector, hereafter called *premierD* (figure 3b). If all of the members in the premier group are defectors, then the division of labour does not start, and the latter group’s players do not have a chance to play the game, so they remain neutral (represented with *).

The equilibria are as follows (figure 3(a) and (b)):

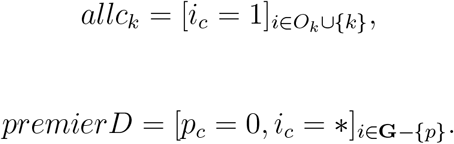

Finally, there is the cooperator-defector mixed equilibrium, when some groups consist of all cooperators and some are all defectors, followed by the neutral groups in *O*_*k*_∪{*k*}. There are two types of mixed equilibrium, when considering from the point of view of *k*; (i) one defector group exists in *U*_*k*_∪{*k*} (figure 3c), or (ii) there are only cooperator groups in *U*_*k*_∪{*k*}, and at least one defector group exists in *D*_*k*_ (figure 3d). This type of mixed equilibrium is hereafter called 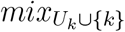. This is represented as follows (figure 3(c)):

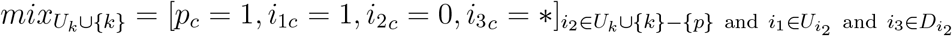

Figure 3(c) shows that *U*_*k*_∪{*k*} is similar to the linear division of labour (Nirjhor and Nakamaru, 2023); if someone chooses defection in a particular group, the task does not get passed to the next group, so the division of labour is stopped, and the later groups’ strategy does not matter.

The second type of mixed equilibrium, (ii), is called 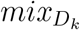. Because of the presence of the branching in *D*_*k*_, we generalize this equilibrium as follows (figure 3(d)):

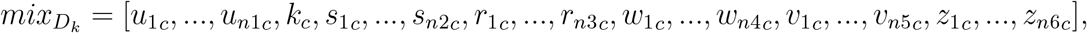

where, *u*_*ic*_ = 1, *k*_*c*_ = 1, *s*_*ic*_ = 1, *r*_*ic*_ = 1, *w*_*ic*_ = 0, *v*_*ic*_ = *, *z*_*ic*_ = 0 where *u*_*i*_ ∈ *U*_*k*_, *s*_*i*_ ∈ *s*_*k*_, *r*_*i*_ ∈ *r*_*k*_, *w*_*i*_ ∈ *w*_*k*_, *v*_*i*_ ∈ *v*_*k*_, and *z*_*i*_ ∈ *z*_*k*_. We define *u* _1_ = *p*. The mutually disjoint sets *s*_*k*_, *r*_*k*_, *w*_*k*_, *v* _*k*_ and *z* _*k*_ are defined as follows,

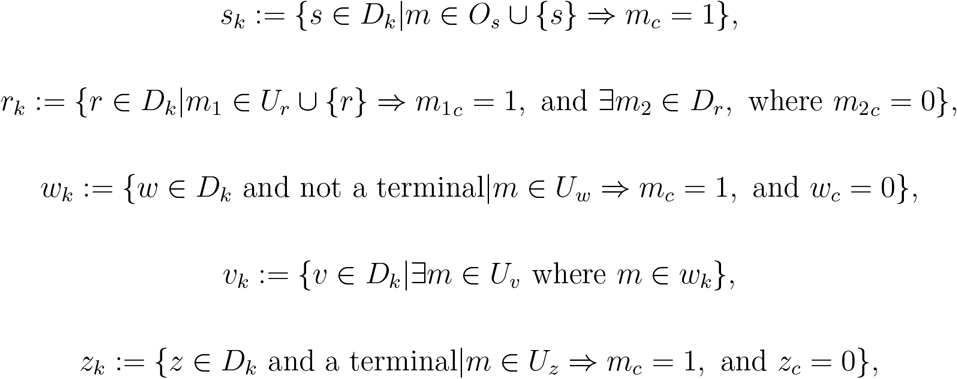

where, |*U*_*k*_| = *n*_1_, |*s*_*k*_| = *n*_2_, |*r*_*k*_| = *n*_3_, |*w*_*k*_| = *n*_4_, |*v*_*k*_| = *n*_5_ and |*z*_*k*_| = *n*_6_. Also,

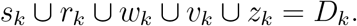

In simple terms, *s*_*k*_ is the set of the groups that are in *D*_*k*_, and their respective trees of the division of labour have a fully cooperative population in all the groups. *r*_*k*_ is the set of the groups in *D*_*k*_, which have fully cooperators, and the upstream groups of their respective trees of the division of labour have fully cooperators. However, at least one group has fully defectors in their downstream. *w*_*k*_ is the set of the groups that are in *D*_*k*_, not terminals, and have a full defector population. *v*_*k*_ is the set of the groups that are in *D*_*k*_ and have a fully defective group in their upstream. The strategies of the members of these groups are neutral, which is represented by * as explained before. *z*_*k*_ is the set of the terminal groups that are in *D*_*k*_, and have a full defector population.

When *k* is the premier, *p*, the 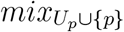 equilibrium does not exist, because there is no *U*_*p*_ and the case of *p* being the defector group is included in the equilibrium *premierD*. When *k* is a terminal, we can consider *O*_*k*_∪{*k*} as a linear division of labour (Nirjhor and Nakamaru, 2023), where 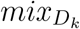 does not exist.

If *k* = *p*, then *O*_*p*_∪{*p*} = **G**, in other words all the groups are in it. If we consider a certain *k*, which is not the premier, eqs. (1)-(3) show that *N*_*k*_ does not have any effect in the dynamics, therefore, it is enough to only consider the stability of the equilibria across *O*_*k*_∪{*k*}.

The local stability of each equilibrium is analyzed (for calculations, refer to Appendix A). Because the equilibria in the general tree graph are hard to write individually, we consider an arbitrary group *k* and define the classes of the equilibria while focusing on that group. The results are mentioned in table A1. To determine the stability of the whole system, we need to consider each of the groups individually and obtain the locally stable conditions of the respective equilibria of those individual focus groups and finally combine their conditions to obtain the locally stable state of the whole system. We shall explain this in detail in the following section using a specific example.

To study a social dilemma situation in the baseline, we consider *g*_*i*_ < *x*_*i*_ for all group *i*. Moreover, if *g*_*i*_ is high enough, it is natural that cooperation among all groups can evolve, and then we do not consider *g*_*i*_ > *x*_*i*_. With this social dilemma condition we summarize the results in table 3, which shows that *premierD* is the only stable equilibrium in the Baseline, because *g*_*p*_ − (1 − *c*_*op*_)*g*_*op*_ < *x*_*p*_ is always held (table A1).

## 3 Two sanction systems

We introduce two types of sanction systems following Nirjhor and Nakamaru (2023). One is called the defector sanction system, where the exact defector is caught and sanctioned with the amount *f*. The finding probability of the exact defector is *ρ*. The other is called the premier sanction system, where if the defection is present, whoever defects, always the player in the premier group is sanctioned with the amount *f*. We study the evolution of cooperation in the system without punishment named the baseline system and then compare its result with the two systems with sanction. The two sanction systems are also compared with each other, to find their effectiveness.

Appendices B and C show the replicator equations for asymmetric games and the results of the local stability analysis of the defector sanction system and the premier sanction system, respectively. Table 4 and A1 summarize the local stability condition for each of the equilibrium points.

**Table 4:**
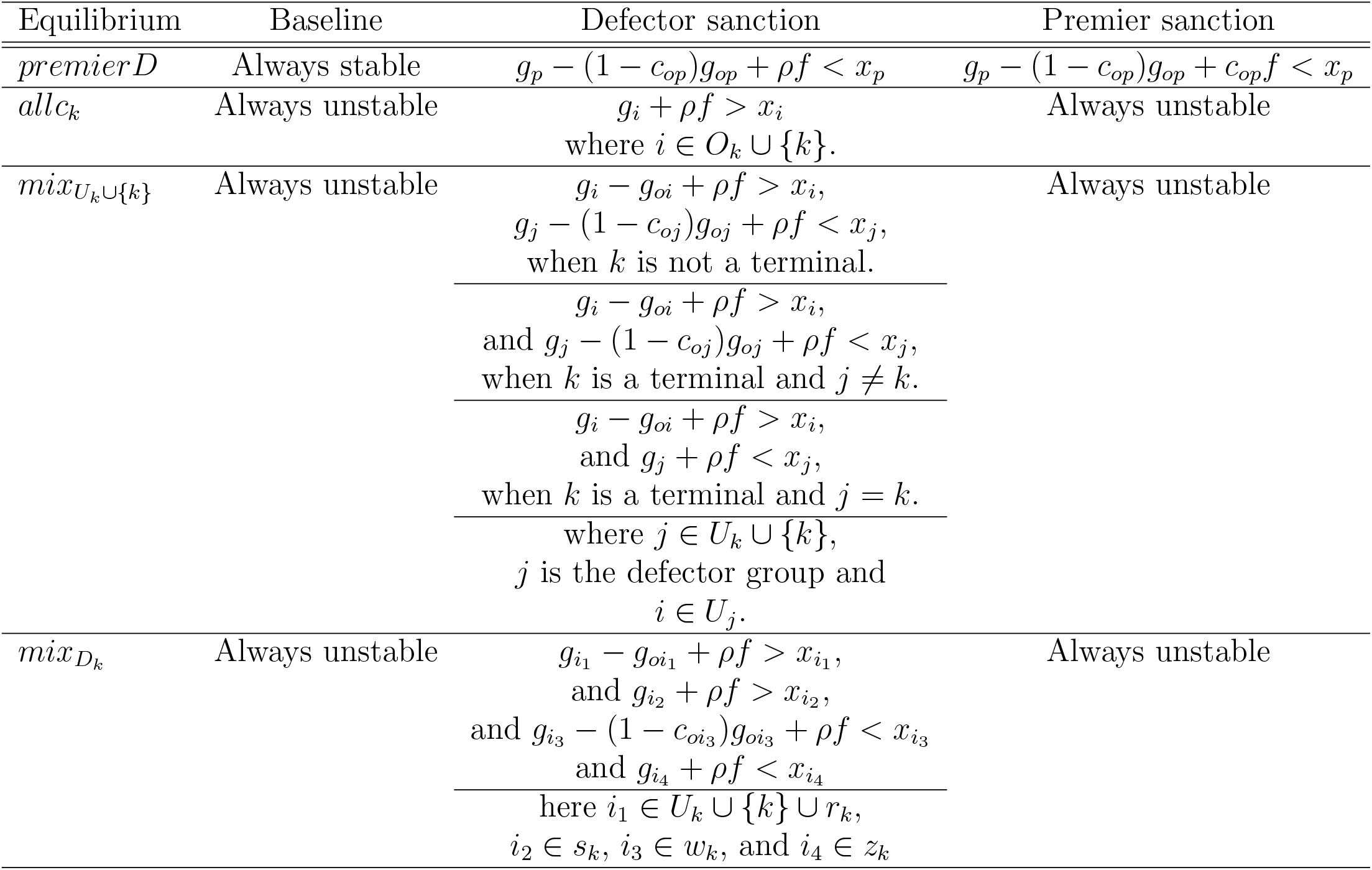
Local stability conditions when *k* is the focal group in a social dilemma

| Equilibrium | Baseline | Defector sanction | Premier sanction |
| --- | --- | --- | --- |
| $premierD$ | Always stable | $g_p - (1 - c_{op})g_{op} + \rho f < x_p$ | $g_p - (1 - c_{op})g_{op} + c_{op}f < x_p$ |
| $allc_k$ | Always unstable | $g_i + \rho f > x_i$<br>where $i \in O_k \cup \{k\}$. | Always unstable |
| $mix_{U_k \cup \{k\}}$ | Always unstable | $g_i - g_{oi} + \rho f > x_i,$<br>$g_j - (1 - c_{oj})g_{oj} + \rho f < x_j,$<br>when $k$ is not a terminal.<br>$g_i - g_{oi} + \rho f > x_i,$<br>and $g_j - (1 - c_{oj})g_{oj} + \rho f < x_j,$<br>when $k$ is a terminal and $j \neq k$.<br>$g_i - g_{oi} + \rho f > x_i,$<br>and $g_j + \rho f < x_j,$<br>when $k$ is a terminal and $j = k$.<br>where $j \in U_k \cup \{k\},$<br>$j$ is the defector group and<br>$i \in U_j.$ | Always unstable |
| $mix_{D_k}$ | Always unstable | $g_{i_1} - g_{oi_1} + \rho f > x_{i_1},$<br>and $g_{i_2} + \rho f > x_{i_2},$<br>and $g_{i_3} - (1 - c_{oi_3})g_{oi_3} + \rho f < x_{i_3}$<br>and $g_{i_4} + \rho f < x_{i_4}$<br>here $i_1 \in U_k \cup \{k\} \cup r_k,$<br>$i_2 \in s_k, i_3 \in w_k,$ and $i_4 \in z_k$ | Always unstable |

We find that *premierD* is the only stable equilibrium in the premier sanction system as well as in the Baseline (table 4 and A1), although the condition is somehow less strict for *allc* to be stable because of the punishment, the same conclusion can be drawn here as well as the baseline. In the defector sanction system, however, all the equilibria are conditionally stable.

We would like to explain the equilibrium using figures 2 and 4. When a defector-sanction system is applied, Figures 2 and 4 are the image of the equilibrium. To obtain the local stability condition for this whole system, **G**, we do the local stability analysis for the equilibrium point with respect to *k* without considering the effect of *N*_*k*_, using eq.(1)-(3). In the case of figures 2 and 4, where **G** converges to the 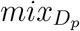 equilibrium, we can obtain the local stability condition of the 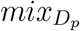 equilibrium in the whole system **G**. Additionally, the 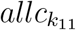 equilibrium with respect to *k*_11_ should be locally stable, the 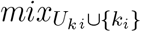 equilibrium with respect to *k*_*i*_ where *i* is 2, 12, 21, 22, and the 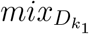 equilibrium with respect to *k*_1_ should be locally stable (figures 2 and 4). The stable equilibria from the perspective of different groups are included in table 2.

**Figure 4:**
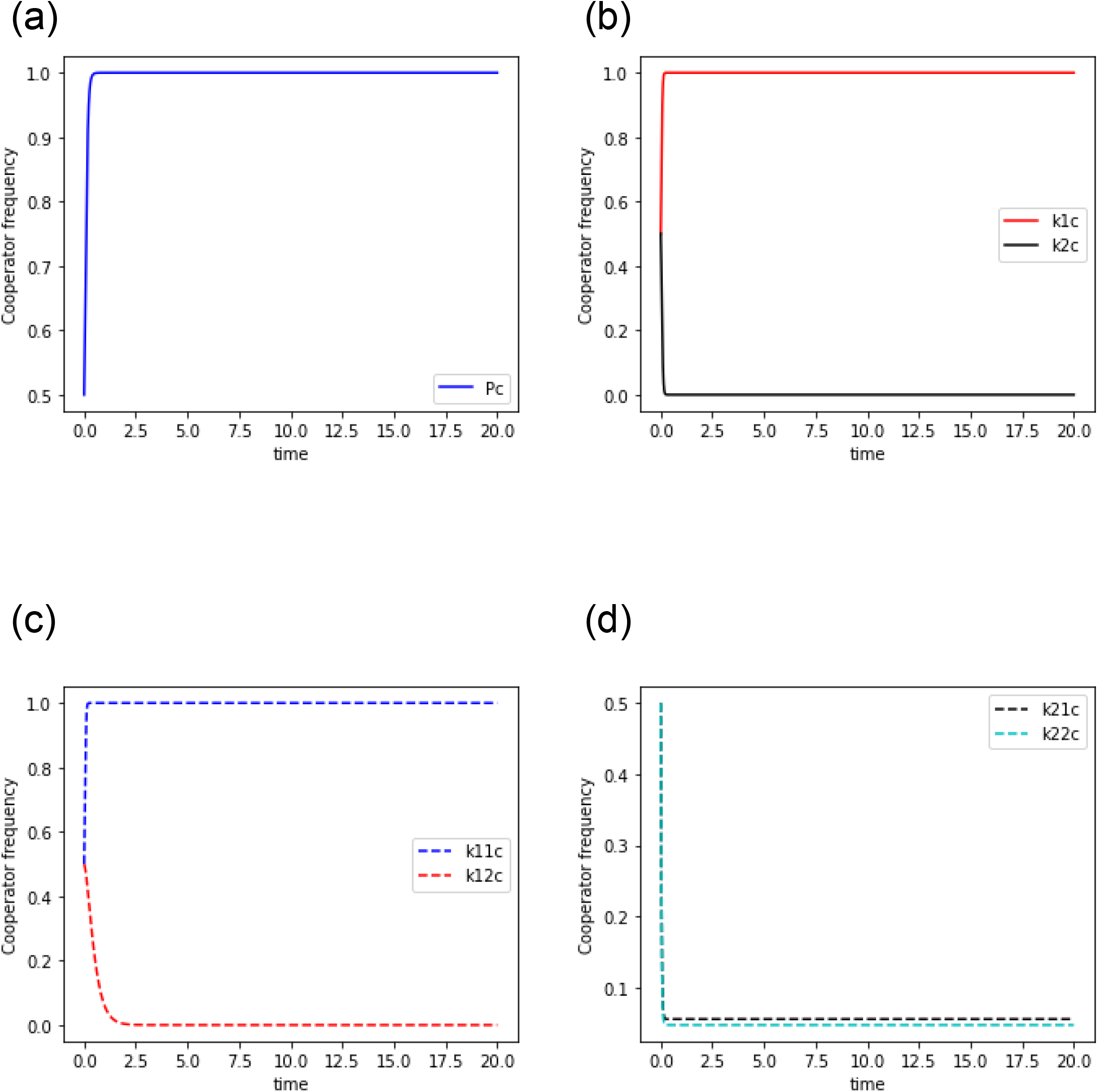
The dynamics of a case as shown in figure 2 in the defector sanction system. 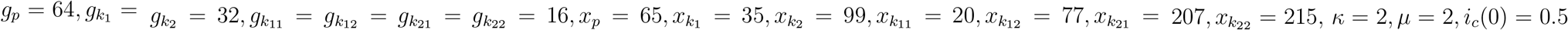 and *ρf* = 58. A point to be noted, this is just an example to show the equilibria according to the relative position of the focal group.

In figure 4, as *x*_*p*_ − *g*_*p*_ + (1 − *c*_*op*_)*g*_*op*_ = 49 < 58 = *ρf*, the premier group becomes full cooperator. As 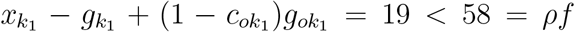 and 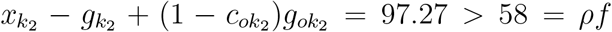, *k*_1_ becomes full cooperator, and *k*_2_ becomes full defector, which in return makes *k*_21_ and *k*_22_ neutral. 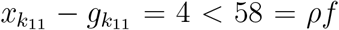 and 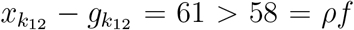, which means *k*_11_ becomes full cooperator and *k*_12_ becomes full defector (table 4). For this reason, with respect to group *k*_11_, the groups in 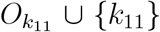 converges to 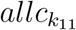. With respect to *k*_12_, the groups in 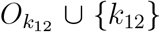 converge to the 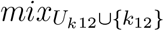 equilibrium and are not influenced by the groups in 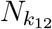.

To understand the dynamics more concretely we do some simplification and numerical analysis in the following section.

## 4 A special case: *κ*-regular, *µ*-branched directed finite tree graph

A *κ*-regular, *µ*-branched directed finite graph is assumed, for making numerical analysis. When *j*s are the immediate branching of *k, g*_*k*_ = Σ_*j*_ *g*_*j*_. In this case, we also consider the distribution of the loss is uniform in each branching. If *k* is at the *µ*_*k*_th branch (where 0 ≤ *µ*_*k*_ ≤ *µ*), we assume, 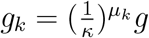.

For the 2-regular, 2-branched graph, the expected values of *g*_*op*_, 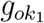, and 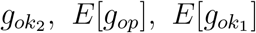 and 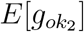, are as follows (figure 2): 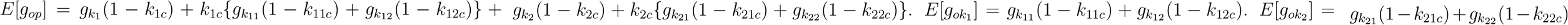, where *k*_22*c*_, for example, is the frequency of cooperators in group *k*_22_ (table 3).

Figure 5 shows the effect of *ρf* on the dynamics in the 2-regular 2-branched directed finite graph. For simplicity, we consider the cost of the cooperation for the groups which are present in the same level branching to be the same.

**Figure 5:**
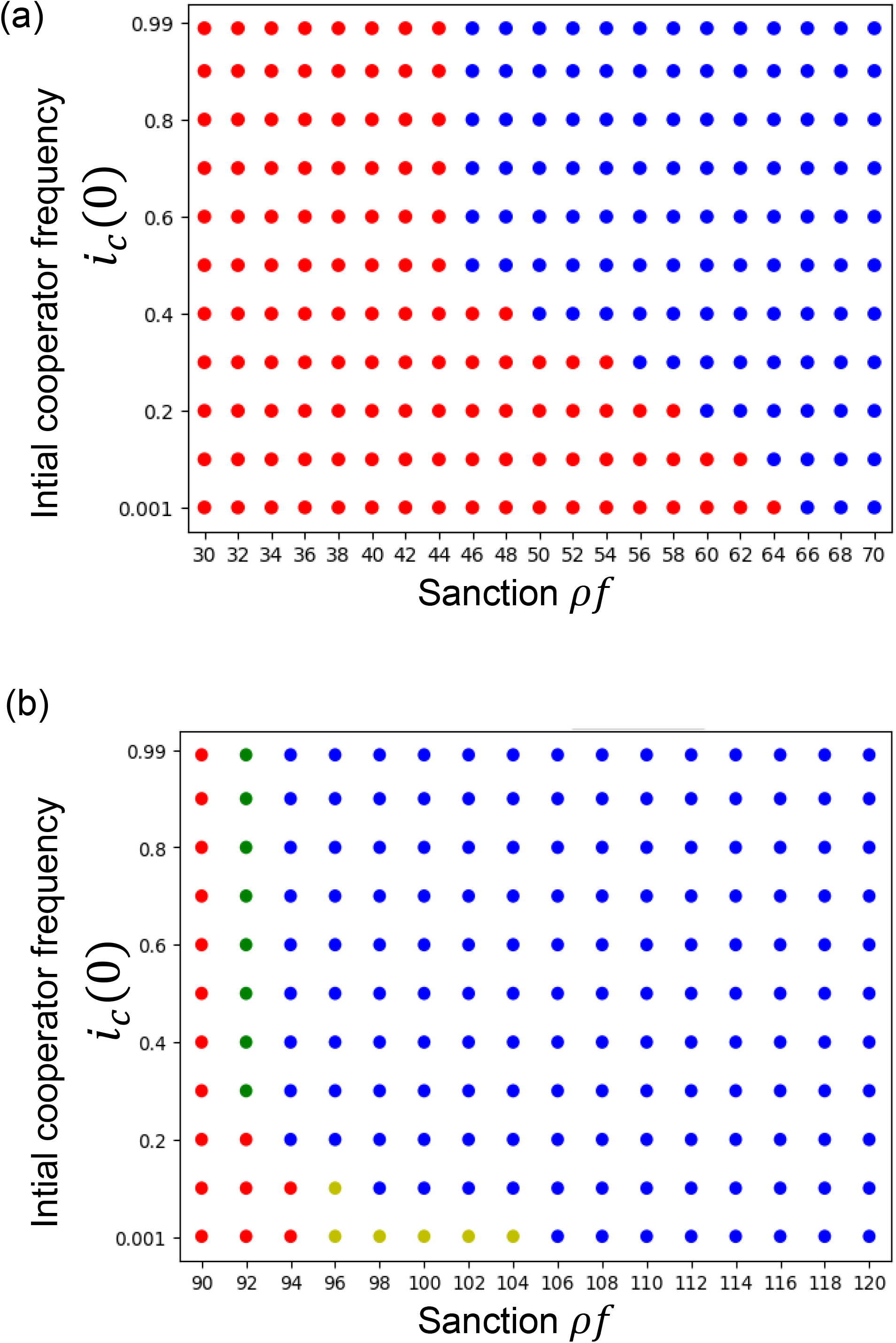
(a) Evolutionary dynamics when cost of the cooperation decreases downstream. 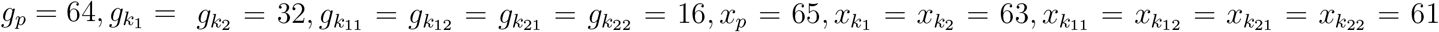. (b) Evolutionary dynamics when cost of the cooperation increases downstream. 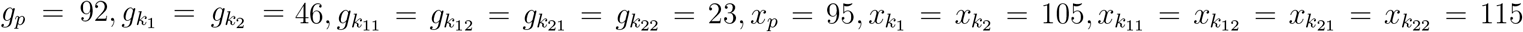. The red dots represent the simulation dynamics converged to the *premierD* equilibrium, the yellow dot represents the mixed equilibrium when the premier group is full cooperator and the groups in the first branching or *k*_1_ and *k*_2_ are full defectors, the green dot represents the mixed equilibrium when the groups premier, *k*_1_ and *k*_2_ are full cooperators and the groups in the 2nd branching *k*_11_, *k*_12_, *k*_21_ and *k*_22_ are full defectors, the blue dot represents when the system moves into *allc*_*p*_ equilibrium, or everyone in every group is a cooperator.

Figure 5(a) is the results of the numerical analysis of equations (B1)-(B3) in Appendix B when the cost of cooperation decreases downstream under the defector sanction system. Figure 5(a) shows that our numerical simulation results match our theoretical prediction (table 4). In figure 5(a), the simulation dynamics converges to the *allc*_*p*_ equilibrium regardless of *i*_*c*_(0) in *ρf* ≥ 66, which matches *ρf* + *g*_*p*_ − (1 − *c*_*op*_)*g*_*op*_ > *x*_*p*_ and *x*_*i*_ < *ρf* + *g*_*i*_ for all the group *i*s. The initial frequencies determine which dynamics converge to the *premierD* or *allc*_*p*_ equilibria, in *ρf* between 46 and 64. This condition matches *x*_*i*_ < *ρf* + *g*_*i*_ for all the group *i*s, however, sometimes matches *ρf* + *g*_*p*_ − (1 − *c*_*op*_)*g*_*op*_ < *x*_*p*_, and sometimes matches *ρf* + *g*_*p*_ − (1 − *c*_*op*_)*g*_*op*_ > *x*_*p*_ depending on the initial frequency of cooperators. The simulation dynamics converge to the *premierD* in *ρf* ≤ 44 which matches *ρf* + *g*_*p*_ − (1 − *c*_*op*_)*g*_*op*_ < *x*_*p*_, regardless of the initial frequencies of cooperators in groups (table 4).

From table 4, it is theoretically clear that the mixed equilibria can also be stable when the cost of the cooperation decreases downstream. This is because *g*_*i*_ − *g*_*oi*_ + *ρf* > *x*_*i*_ > *x*_*j*_ > *g*_*j*_ − (1 − *c*_*oj*_)*g*_*oj*_ + *ρf* as well as *g*_*i*_ − *g*_*oi*_ + *ρf* > *x*_*i*_ > *x*_*j*_ > *g*_*j*_ + *ρf* have no contradiction when *j* ∈ *D*_*i*_ (table 4). However, as *c*_*ok*_ and *g*_*ok*_ are hard to predict, it is hard to predict when the mixed equilibria might be stable.

Figure 5(b) shows the numerical simulation outcomes when the cost of the cooperation increases downstream. The simulation dynamics converges to the *premierD* in *ρf* ≤ 90 which matches *ρf* + *g*_*p*_ − (1 − *c*_*op*_)*g*_*op*_ < *x*_*p*_, regardless of *i*_*c*_(0) (table 4). When *ρf* is 92 in figure 5(b), there is a co-presence of green dots and red dots, which indicates the bistability predicted by table 4; the initial frequency of cooperators in groups determines if the dynamics converge to either the mixed equilibrium 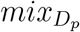 or the *premierD*. When *ρf* + *g*_*p*_ − *g*_*op*_ > *x*_*p*_, and *ρf* + *g*_*i*_ − (1 − *c*_*oi*_)*g*_*oi*_ < *x*_*i*_ holds for some group *i*, it is 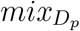, and when *ρf* + *g*_*p*_ − *g*_*op*_ < *x*_*p*_ it is *premierD*. For this reason the 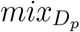 and *premierD* can be bistable. The red dots and the blue dots are co-present when *ρf* is 94 in figure 5(b), which matches the theoretical prediction; the bistability between the equilibria *premierD* and *allc*_*p*_. When *ρf* + *g*_*p*_ − (1 − *c*_*op*_)*g*_*op*_ > *x*_*p*_ and *x*_*i*_ < *ρf* + *g*_*i*_ for all group *i*s it is *allc*_*p*_, and when *ρf* + *g*_*p*_ − (1 − *c*_*op*_)*g*_*op*_ < *x*_*p*_ it is *premierD* (table 4). It is shown with the co-presence of blue and yellow dots when *ρf* is from 96 to 104 in figure 5(b), which matches table 4; the bistability between the 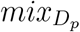 and *allc*_*p*_ is found when the conditions *ρf* + *g*_*i*_ > *x*_*i*_ for all the group *i*s and *ρf* + *g*_*j*_ − (1 − *c*_*oj*_)*g*_*oj*_ < *x*_*j*_ for some group *j* which is not a terminal, are both present depending on the initial condition along with *ρf* + *g*_*p*_ − (1 − *c*_*op*_)*g*_*op*_ < *x*_*p*_ (table 4). However, the mixed equilibrium 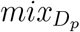, with a terminal group being the defector group shown in the green dots can not be bistable with *allc*_*p*_ shown in blue dots, because their stability conditions contradict each other (table 4). When *ρf* ≥ 106, the simulation dynamics converge to *allc*_*p*_, which is also predicted by table 4.

## 5 Discussion and conclusion

We took a model of the division of labour in a finite tree graph and studied the effect of sanctions on it. There is a premier group and then the division of labour is branched from it. Each node of the tree graph has a group which has a role in the division of labour. The task flows from the upstream to the downstream and gets divided through the branching. If a player who is randomly selected from a group chooses defection, the division of labour stops there and everyone in every group needs to bear a loss according to their position. We compare the evolution of cooperation in the baseline system (which has no sanction) with the two sanction systems named the defector sanction system and the premier sanction system. We studied the general model and found three equilibria when the defector sanction system is applied in the social dilemma situation, 1) *premierD*, where all players in the whole premier group choose defection, 2) *allc* where everyone in every group is a cooperator, and 3) mixed equilibrium, where the premier group consists of only cooperators, some other groups also are full of cooperators, and somewhere in the network there are one or more group/groups who have a whole population of defectors. We did the local stability analysis of these equilibria. Then, for doing numerical analysis, we considered a special case and verified the results of the general case.

The previous theoretical studies show that the cooperation evolves in a network structured population when *b/c* > *k*, where each node representing a single player has *k* regular links and *b* is the benefit from a cooperator and *c* is a cost of cooperation (e.g. Nowak, 2006). However, in our study, the benefit to a player given by a cooperator from the upstream group is cancelled out in equations and does not have any effect on the evolution of cooperation. This means that our results cannot be summarized by the ratio of *b* and *c*. This result is the same as the linear division of labour (Nirjhor and Nakamaru, 2023). This is because both the cooperators and defectors in a group receive the benefit from the cooperation in the upstream groups regardless of their own strategy. This can be interpreted as the salary given to an employee in this sort of division of labour has no effect on the evolution of cooperation.

The loss via defection becomes distributed in the branches as *g*_*k*_ = Σ*j* are immediate branching of *k g*_*j*_. The loss via defection *g*_*k*_ is subjective to the task assigned to the group *k*. Because of this setting a group *k*’s evolutionary dynamics are only affected by the action of groups in *O*_*k*_∪{*k*}, as those are the groups directly associated with the action that is assigned to *k*. If there is a defection in the upstream of *k*, the loss to *k* is the same as if the task is not being completed by a player in group *k*. A defection downstream of *k* means a part of the task assigned to group *k* is not eventually fully completed, which in turn affects the payoff of *k* as *g*_*ok*_. However, because of this setting the tasks assigned to groups present in *N*_*k*_ have no relation with the task of group *k*, and because of that, their defection does not affect the evolutionary dynamics of *k*. That is why *g*_*nk*_ is canceled out from the replicator equation for the dynamics of group *k*. This means that a group’s decisions are influenced only by that part of the network with which the nodes have a direct hierarchical connection with that particular group. In simpler terms, a group is influenced by other groups which are either in its hierarchical upstream or downstream, not the groups which are branched separately from its upstream but belong to the same network. This result can be applied to the division of labour of the government, as it shows that a corrupt/honest sector can exist independently and in a government, even when other sectors of the government are honest/corrupt.

With comparison with the linear division of labour (Nirjhor and Nakamaru, 2023), we find that the mixed equilibria can be stable in the baseline as well as the premier sanction system, when we do not consider the social dilemma situation, in other words, *g*_*i*_ < *x*_*i*_ for all the group *i*s does not necessarily hold. In Nirjhor and Nakamaru (2023), the mixed equilibria were unstable in the baseline and the first role sanction system regardless of social dilemma or not. We theoretically find that the mixed equilibrium can be stable even when the cost is decreasing downstream in the defector sanction system. In the linear division of labour of Nirjhor and Nakamaru (2023), the mixed equilibrium is never stable when the cost is decreasing downstream.

We should mention the applicability of this study to the supply chain. Our study can be applied to the multilayered subcontract (e.g. Tam et al, 2011) which has the tree structure assumed in our study. There are various networks among roles and stakeholders in the supply chain. In Lambert and Cooper (2000), the generalized supply chain network was shown to be an uprooted tree-like, where there is a central body that can be considered as the stem of the tree from which branches spread in both directions of the root and the shoot. From one direction the branches merge upstream towards the central body showing many divisions of the labour merging into the completion of a single labour, and from there the branches split downstream showing the labour is being divided. Our study addresses the evolution of cooperation in the later part of the supply chain where the labour is being divided downstream. When each player is assumed to be located at each node of trees and to interact with the neighbors, the effect of the merge of networks or directed cycles on the evolution of cooperation has been investigated (e.g. Su et al., 2022). As we assume that a group is located at each node of trees, our model and results would be different from the previous studies. In our future research, we wish to address the problem of the evolution of cooperation in the former part of the supply chain as well, where the division of labour merges together upstream to complete a single labour, and then extend our study to the uprooted tree-like networks.

## Supporting information

Code for figures 4 and 5

## Acknowledgement

NMSA was supported by JST, the establishment of university fellowships towards the creation of science technology innovation, Grant Number JPMJFS2112. MN was supported by JSPS KAKENHI Grant Number JP21K01626.

## Appendix A The local stability analysis of the baseline model

On the basis of Eqs. 1-3 in the main text, we make the time differential equations for *O*_*k*_∪{*k*}. Then, we calculate Jacobian matrix from the differential equations for analyzing the local stability of the equilibria based on *O*_*k*_∪{*k*}. All Jacobian matrices are lower triangular matrices here, so the eigenvalues are just the main diagonal entries.

The Jacobian matrix for *premierD* is *J*(*premierD*). If *ξ* is a main diagonal component of *J*(*premierD*) then,

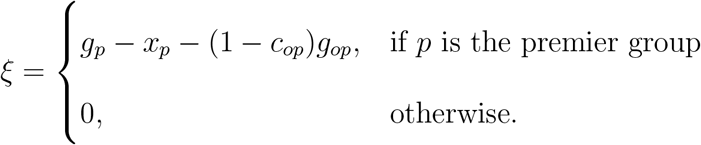

When *g*_*p*_ − (1 − *c*_*op*_)*g*_*op*_ < *x*_*p*_, *PremierD* is locally stable. As *g*_*p*_, *x*_*p*_, *g*_*op*_ > 0, 0 ≤ *c*_*op*_ ≤ 1, *g*_*op*_ ≤ *g*_*p*_, and *g*_*p*_ is always lower than *x*_*p*_ in the social dilemma. Therefore, *PremierD* is always locally stable.

The Jacobian matrix for *allc*_*k*_ is *J*(*allc*_*k*_). If *ξ* is a main diagonal component of *J*(*allc*_*k*_) then

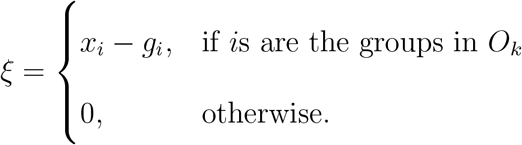

As the baseline model has the social dilemma situation, *x*_*i*_ > *g*_*i*_. Then, *allc*_*k*_ is not locally stable.

The Jacobian matrix for 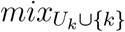 is 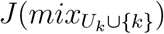. If *ξ* is a main diagonal component of 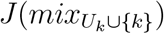 and *k* is not a terminal, or *k* is a terminal however, *k* is not the defector group, then

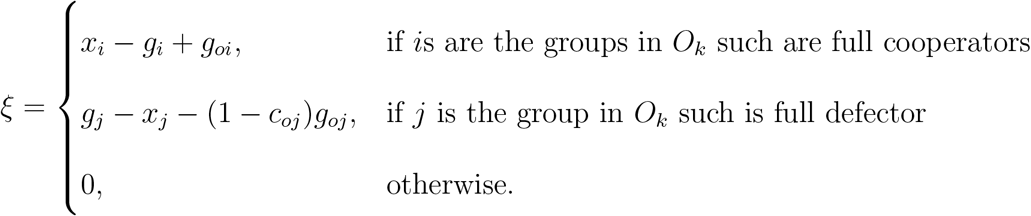

If *ξ* is a main diagonal component of 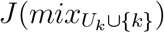 and *k* is a terminal and *k* is the defector group, then

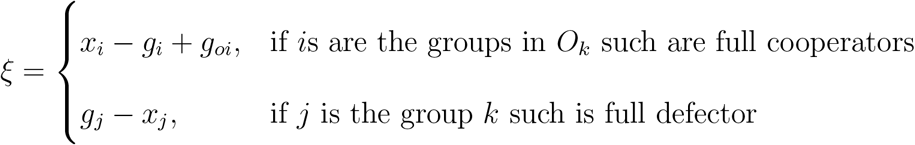

As the baseline model has the social dilemma situation, *x*_*i*_ > *g*_*i*_ − *g*_*oi*_. Therefore, the 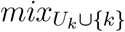 is not stable.

The Jacobian matrix for 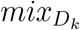 is 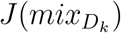. If *ξ* is a main diagonal component of 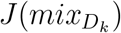 then

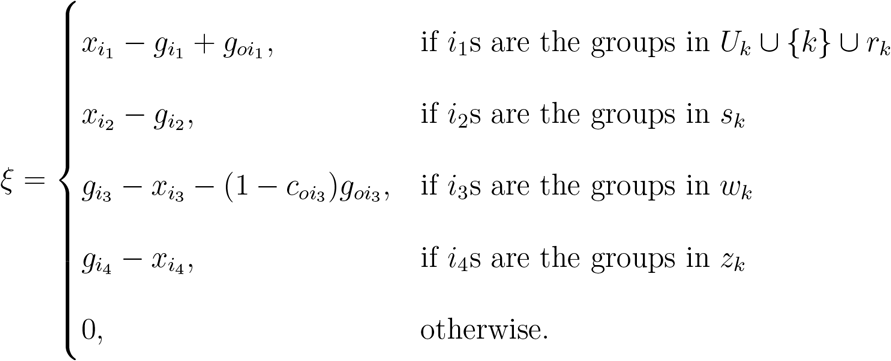

This equilibrium is always unstable as the baseline model has the social dilemma situation, 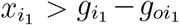, because for 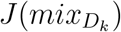, *U*_*k*_ ∪{*k*} is never an empty set. Table A1 summarizes the local stability condition of four equilibrium points in the baseline model.

**Table A1:**
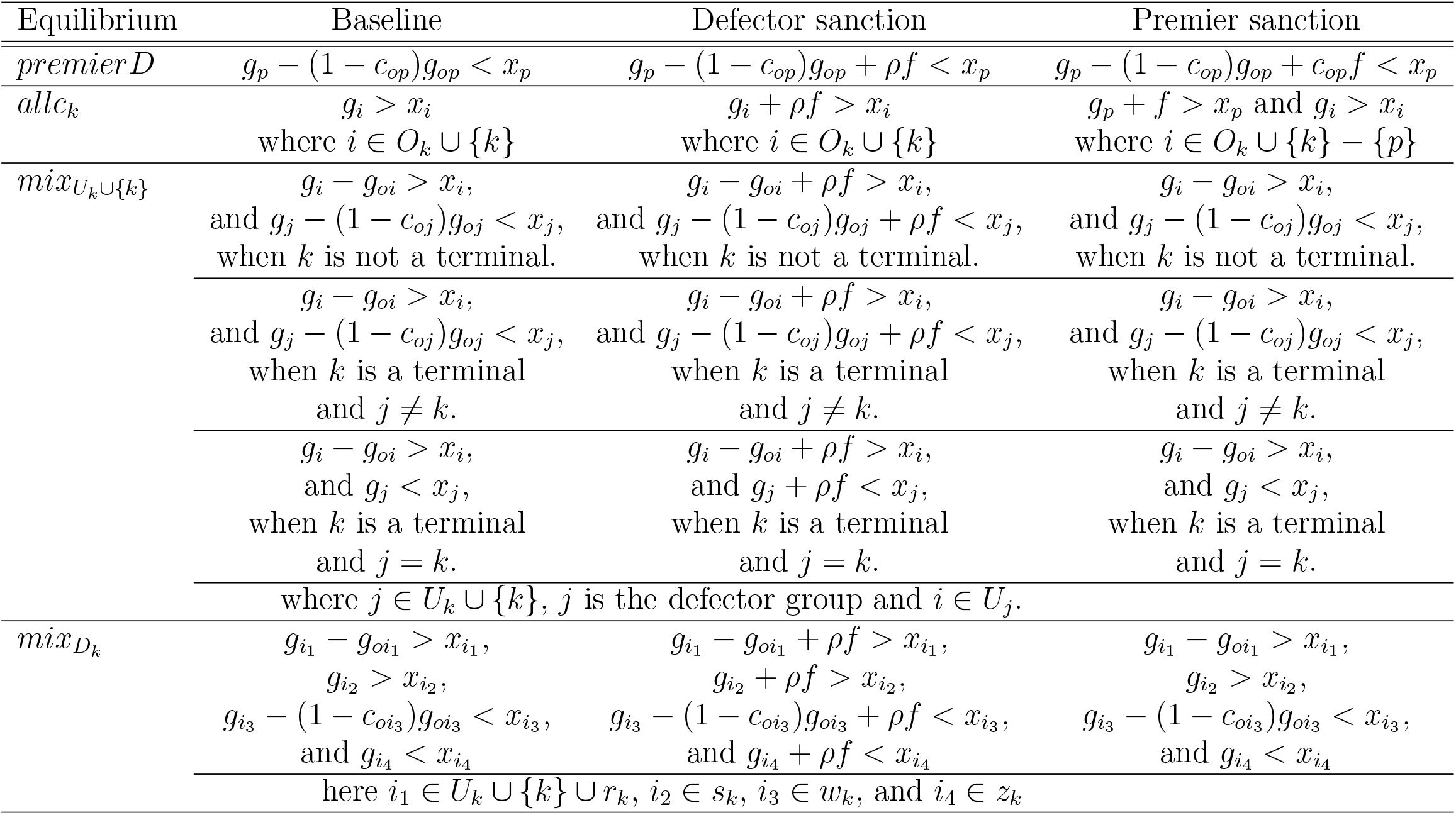
Local stability conditions when *k* is the focal group

## Appendix B The defector sanction system

The payoff matrices of the defector sanction system are shown in tables B1a, B1b, and B1c. The payoffs are the same as the baseline except while choosing defection, the players are subjected to the sanction *ρf*. Premier’s payoff in the defector sanction system when he is a cooperator, Π_*cp*_ is *b*_*p*_ − *x*_*p*_ − *g*_*op*_ + *c*_*op*_*g*_*op*_. Premier’s payoff in the defector sanction system when he is a defector, Π_*dp*_, is *b*_*p*_ − *g*_*p*_ − *ρf*. Therefore, the replicator equation is,

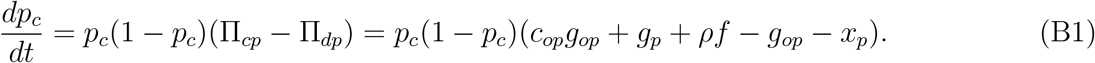

Any player *k*’s payoff when *k* is neither premier nor a terminal, and *k* is a cooperator, Π_*ck*_, is *c*_*ok*_(*b*_*k*_ − *x*_*k*_) − *d*_*uk*_*g*_*k*_ + *d*_*dk*_(*b*_*k*_ − *x*_*k*_ − *g*_*ok*_) − *g*_*nk*_. When *k* is a defector the payoff, Π_*dk*_, is *c*_*ok*_(*b*_*k*_ − *g*_*k*_ − *ρf*) − *d*_*uk*_*g*_*k*_ + *d*_*dk*_(*b*_*k*_ − *g*_*k*_ − *ρf*) − *g*_*nk*_. Therefore, the replicator equation is,

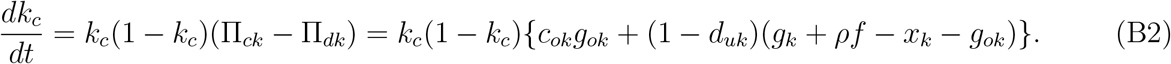

Any player *n*’s payoff when *n* is a terminal, and *n* is a cooperator, Π_*cn*_, is *c*_*on*_(*b*_*n*_ − *x*_*n*_) − *d*_*un*_*g*_*n*_ − *g*_*nn*_. When *n* is a defector the payoff, Π_*dn*_, is, *c*_*on*_(*b*_*n*_ − *g*_*n*_ − *ρf*) − *d*_*un*_*g*_*n*_ − *g*_*nn*_. Therefore, the replicator equation is,

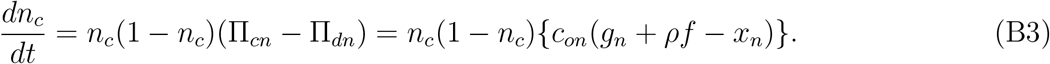

Here, the benefit *b*_*k*_ given by a cooperator of the upstream as well as the term *g*_*nk*_ which represents the relation of *k*s dynamics with *N*_*k*_ are both canceled in the replicator equation. Therefore, they do not have any effect on the dynamics.

On the basis of Eqs. B1-B3, we make the time differential equations for *O*_*k*_∪{*k*}. Then, we calculate Jacobian matrix from the differential equations for analyzing the local stability of the equilibria based on *O*_*k*_∪{*k*}.

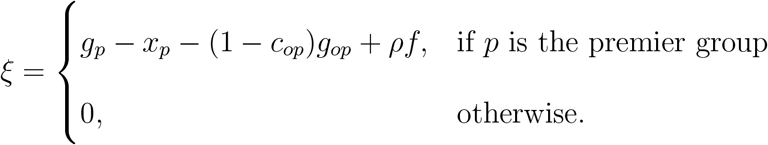

When *g*_*p*_ − (1 − *c*_*op*_)*g*_*op*_ + *ρf* < *x*_*p*_, *PremierD* is locally stable.

In the defector sanction system the Jacobian matrix for *allc*_*k*_ is *J*(*allc*_*k*_). If *ξ* is a main diagonal component of *J*(*allc*_*k*_) then

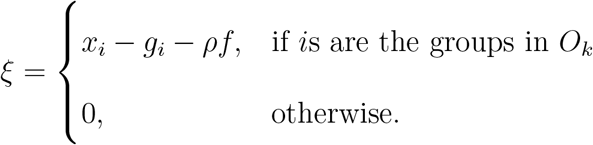

When *x*_*i*_ < *g*_*i*_ + *ρf*, for all *i*s, *allc*_*k*_ is stable.

The Jacobian matrix for 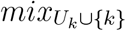 is 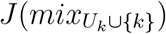. If *ξ* is a main diagonal component of 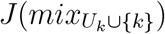 and *k* is not a terminal, or *k* is a terminal however, *k* is not the defector group, then

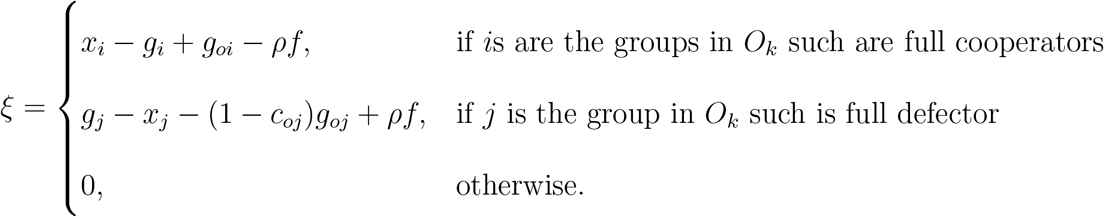

When *x*_*i*_ < *g*_*i*_ − *g*_*oi*_ + *ρf* for all *i*s, and *g*_*j*_ − (1 − *c*_*oj*_)*g*_*oj*_ + *ρf* < *x*_*j*_, then here the 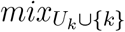 is stable. If *ξ* is a main diagonal component of 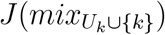 and *k* is a terminal and *k* is the defector group, then

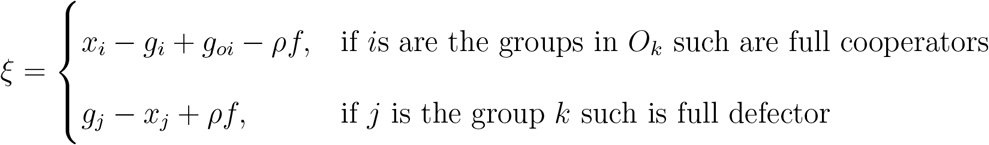

The Jacobian matrix for 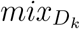 is 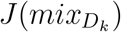. If *ξ* is a main diagonal component of 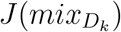 then

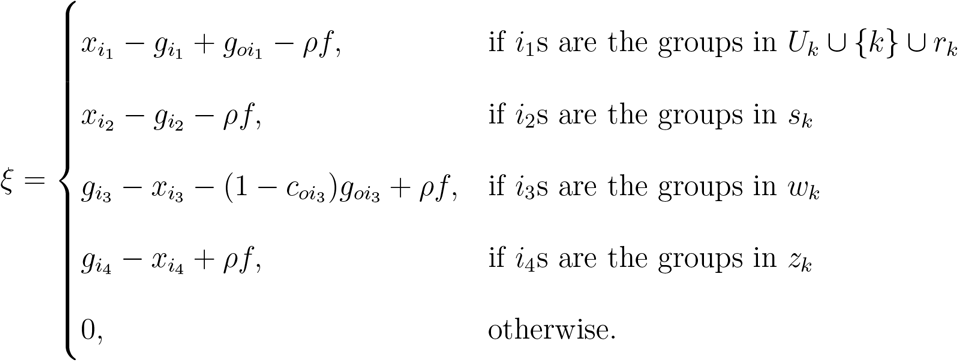

When 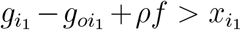 and 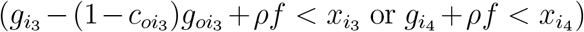 and/or 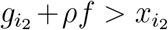, then the 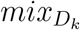 is stable. Table A1 summarizes the local stability condition of four equilibrium points in the defector sanction system.

**Table B1a:**
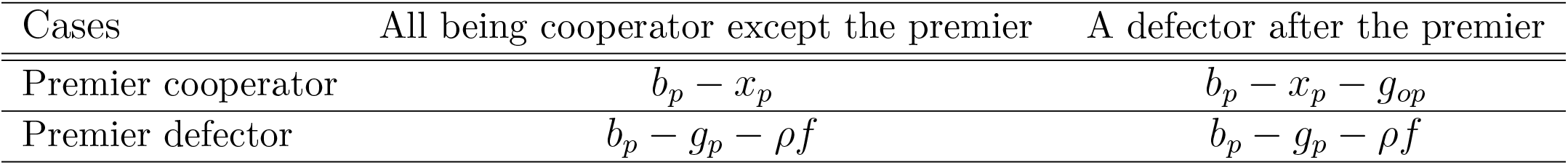
The payoff matrix in Defector sanction system for Premier

**Table B1b:** The payoff matrix in the defector sanction system for *k* ≠ *p*

| Cases | All being cooperator in $\mathbf{O}_k$ | A defector in $U_k$ | A defector in $D_k$ |
| --- | --- | --- | --- |
| $k$ cooperator | $b_k - x_k - g_{nk}$ | $-g_k - g_{nk}$ | $b_k - x_k - g_{ok} - g_{nk}$ |
| $k$ defector | $b_k - g_k - g_{nk} - \rho f$ | $-g_k - g_{nk}$ | $b_k - g_k - g_{nk} - \rho f$ |

**Table B1c:** The payoff matrix in the Defector sanction system for *n* in terminal

| Cases | All being cooperator in $\mathbf{O}_n$ | A defector in $U_n$ |
| --- | --- | --- |
| $n$ cooperator | $b_n - x_n - g_{nn}$ | $-g_n - g_{nn}$ |
| $n$ defector | $b_n - g_n - g_{nn} - \rho f$ | $-g_n - g_{nn}$ |

## Appendix C The premier sanction system

The payoff matrices of the premier sanction system are shown in table C1a, C1b, and C1c. The payoffs are the same as the baseline except while there is a defection in downstream the player in the premier group is subjected to the sanction *f*. Premier’s payoff in the premier sanction system when he is a cooperator, Π_*cp*_, is *b*_*p*_ − *x*_*p*_ − (1 − *c*_*op*_)(*g*_*op*_ + *f*). Premier’s payoff in the premier sanction system when he is a defector, Π_*dp*_, is *b*_*p*_ − *g*_*p*_ − *f*. Therefore, the replicator equation is,

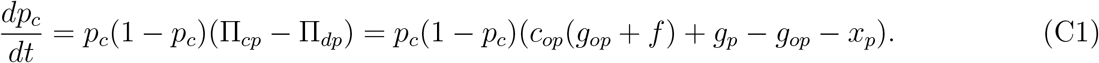

Any player *k*’s payoff when *k* is neither premier nor a terminal, and *k* is a cooperator, Π_*ck*_, is *c*_*ok*_(*b*_*k*_ − *x*_*k*_) − *d*_*uk*_*g*_*k*_ + *d*_*dk*_(*b*_*k*_ − *x*_*k*_ − *g*_*ok*_) − *g*_*nk*_. When *k* is a defector the payoff Π_*dk*_, is *c*_*ok*_(*b*_*k*_ − *g*_*k*_) − *d*_*uk*_*g*_*k*_ + *d*_*dk*_(*b*_*k*_ − *g*_*k*_) − *g*_*nk*_. Therefore, the replicator equation is,

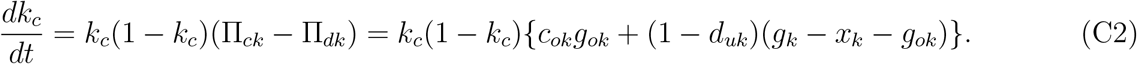

Any player *n*’s payoff when *n* is a terminal, and *n* is a cooperator, Π_*cn*_, is *c*_*on*_(*b*_*n*_ − *x*_*n*_) − *d*_*un*_*g*_*n*_ − *g*_*nn*_. When *n* is a defector the payoff, Π_*dn*_, is *c*_*on*_(*b*_*n*_ − *g*_*n*_) − *d*_*un*_*g*_*n*_ − *g*_*nn*_. Therefore, the replicator equation is,

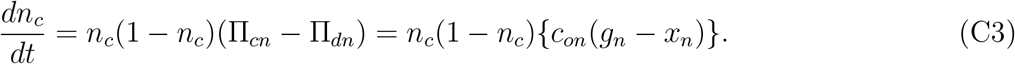

Here, the benefit *b*_*k*_ given by a cooperator of the upstream as well as the term *g*_*nk*_ which represents the relation of *k*s dynamics with *N*_*k*_ are both cancelled in the replicator equation. Therefore, they do not have any effect on the dynamics.

On the basis of Eqs. C1-C3, we make the time differential equations for *O*_*k*_∪{*k*}. Then, we calculate Jacobian matrix from the differential equations for analyzing the local stability of the equilibria based on *O*_*k*_∪{*k*}. The Jacobian matrix for *premierD* is *J*(*premierD*). If *ξ* is a main diagonal component of *J*(*premierD*) then,

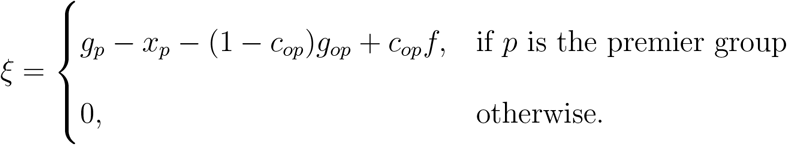

When *g*_*p*_ − (1 − *c*_*op*_)*g*_*op*_ + *c*_*op*_*f* < *x*_*p*_, *PremierD* is locally stable.

In the premier sanction system the Jacobian matrix for *allc*_*k*_ is *J*(*allc*_*k*_). If *ξ* is a main diagonal component of *J*(*allc*_*k*_) then

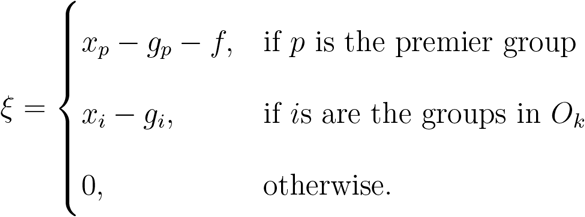

As the model has the social dilemma situation, *x*_*i*_ > *g*_*i*_. Then, *allc*_*k*_ is not locally stable.

The Jacobian matrix for 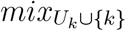 is 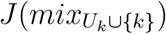. If *ξ* is a main diagonal component of 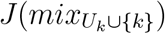 and *k* is not a terminal, or *k* is a terminal however, *k* is not the defector group, then

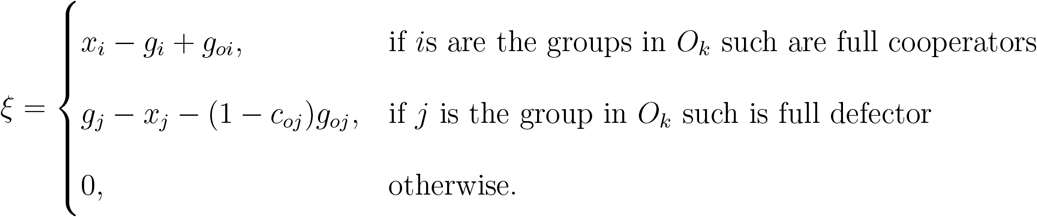

If *ξ* is a main diagonal component of 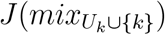 and *k* is a terminal and *k* is the defector group, then

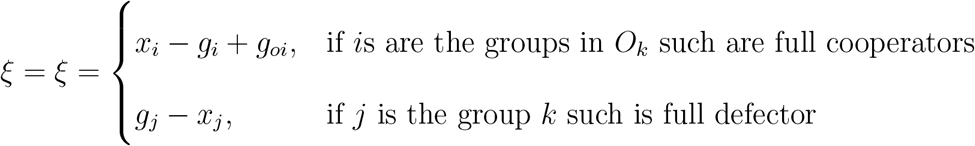

As the model has the social dilemma situation, *x*_*i*_ > *g*_*i*_ − *g*_*oi*_. Therefore, the 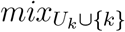 is not stable.

The Jacobian matrix for 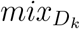 is 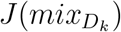. If *ξ* is a main diagonal component of 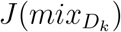 then

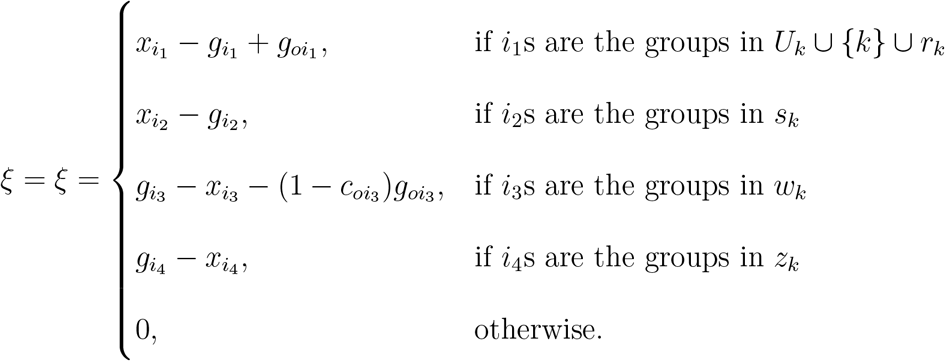

This equilibrium is always unstable as the model has the social dilemma situation, 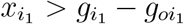, because for 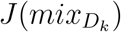, *U*_*k*_∪{*k*} is never an empty set. Table A1 summarizes the local stability condition of four equilibrium points in the premier sanction system.

**Table C1a:**
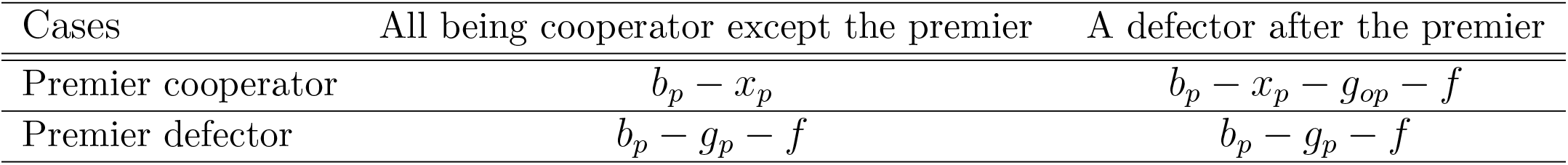
The payoff matrix in Premier sanction system for Premier

**Table C1b:** The payoff matrix in the premier sanction system for *k* ≠ *p*

| Cases | All being cooperator in $\mathbf{O}_k$ | A defector in $U_k$ | A defector in $D_k$ |
| --- | --- | --- | --- |
| $k$ cooperator | $b_k - x_k - g_{nk}$ | $-g_k - g_{nk}$ | $b_k - x_k - g_{ok} - g_{nk}$ |
| $k$ defector | $b_k - g_k - g_{nk}$ | $-g_k - g_{nk}$ | $b_k - g_k - g_{nk}$ |

**Table C1c:** The payoff matrix in the Premier sanction system for *n* in terminal

| Cases | All being cooperator in $\mathbf{O}_n$ | A defector in $U_n$ |
| --- | --- | --- |
| $n$ cooperator | $b_n - x_n - g_{nn}$ | $-g_n - g_{nn}$ |
| $n$ defector | $b_n - g_n - g_{nn}$ | $-g_n - g_{nn}$ |

