## Supplementary material for "The evolution of cooperation in the unidirectional division of labor on a tree network": Code for figures 4 and 5

```
%This is the general MATLAB R2023a code for the figures 4 and 5 of manuscript titled "The evolution of cooperation in the unidirectional division of labor on a tree network"

%Title: Dynamics in the evolution of cooperation in the unidirectional division of labour on a tree network

%Authors: MD Sams Afif Nirjhor and Mayuko Nakamaru

%Institute: Tokyo Institute of Technology

%Date: 14 June, 2023

function main()



    %cost of cooperation for each group

    xp = 65;

    x1 = 63;

    x2 = 63;

    x11 = 61;

    x12 = 61;

    x21 = 61;

    x22 = 61;

    %loss via defection from each group

    gp = 64;

    g1 = 32;

    g2 = 32;

    g11 = 16;

    g12 = 16;

    g21 = 16;

    g22 = 16;

    %preparing the vectors to store the end points of each dynamics

    fP=[];

    fk1=[];
```

```
fk2=[];
fk11=[];
fk12=[];
fk21=[];
fk22=[];
Rhof=[];
ic=[];
%initial condition for the dyanmics
Z0 = [0.99, 0.99, 0.99, 0.99, 0.99, 0.99, 0.99;
     0.9, 0.9, 0.9, 0.9, 0.9, 0.9, 0.9;
     0.8, 0.8, 0.8, 0.8, 0.8, 0.8, 0.8;
     0.7, 0.7, 0.7, 0.7, 0.7, 0.7, 0.7;
     0.6, 0.6, 0.6, 0.6, 0.6, 0.6, 0.6;
     0.5, 0.5, 0.5, 0.5, 0.5, 0.5, 0.5;
     0.4, 0.4, 0.4, 0.4, 0.4, 0.4, 0.4;
     0.3, 0.3, 0.3, 0.3, 0.3, 0.3, 0.3;
     0.2, 0.2, 0.2, 0.2, 0.2, 0.2, 0.2;
     0.1, 0.1, 0.1, 0.1, 0.1, 0.1, 0.1;
     0.001, 0.001, 0.001, 0.001, 0.001, 0.001, 0.001];



for i = 30:2:70
   rhof = i
   for j = 1:1:11
      %solving the replicator equation system
      z0 = Z0(j,:);
```

```
n = 2000;
tspan = linspace(0,30000,n);

[t, Z] = ode45(@(t,z) model(z,t,xp,x1,x2,x11,x12,x21,x22,gp,g1,g2,g11,g12,g21,g22,rhof), tspan,
z0);

P = Z(:,1);
k1 = Z(:,2);
k2 = Z(:,3);
k11 = Z(:,4);
k12 = Z(:,5);
k21 = Z(:,6);
k22 = Z(:,7);
disp(P());
%storing the endpoints of the dynamics for respective amount of sanction and initial condition
fP=[fP,P(end)];
fk1=[fk1,k1(end)];
fk2=[fk2,k2(end)];
fk11=[fk11,k11(end)];
fk12=[fk12,k12(end)];
fk21=[fk21,k21(end)];
fk22=[fk22,k22(end)];
Rhof=[Rhof,rhof]
ic=[ic,z0(1)]

%plotting the results
```

```
            figure;
            plot(t, P, 'b', 'LineWidth', 2);
            hold on;
            plot(t, k1, 'r-', 'LineWidth', 2);
            plot(t, k2, 'm-', 'LineWidth', 2);
            plot(t, k11, 'k--', 'LineWidth', 2);
            plot(t, k12, 'y-', 'LineWidth', 2);
            plot(t, k21, 'k-', 'LineWidth', 2);
            plot(t, k22, 'b-', 'LineWidth', 2);
            xlabel('time');
            ylabel('Cooperator frequency');
            legend('P', 'k1c', 'k2c', 'k11c', 'k12c', 'k21c', 'k22c');
            set(gca, 'FontSize', 14);
            grid off;
        end


    end
%storing the endpoints of each dynamics in an excel sheet, which can latter be plotted for figure 5
DS1 = table(ic', Rhof', fP', fk1', fk2', fk11', fk12', fk21', fk22', ...
            'VariableNames', {'ic','rhof', 'p', 'k1', 'k2', 'k11', 'k12', 'k21', 'k22'});

writetable(DS1, 'outputtree.xlsx', 'Sheet', 1);


end
%defining function for model
function dzdt = model(z,t,xp,x1,x2,x11,x12,x21,x22,gp,g1,g2,g11,g12,g21,g22,rhof)
    %frequency of cooperators in each group
    P = z(1);
```

```
k1 = z(2);
k2 = z(3);
k11 = z(4);
k12 = z(5);
k21 = z(6);
k22 = z(7);
%defining the potential losses
gop = (k1*(k11*(1-k12)*g12+(1-k11)*k12*g11+(1-k11)*(1-k12)*(g11+g12))+(1-k1)*g1+(1-k2)*g2+
k2*(k21*(1-k22)*g22+(1-k21)*k22*g21+(1-k21)*(1-k22)*(g21+g22)));
go1 = ((1-k11)*g11+(1-k12)*g12);
go2 = ((1-k21)*g21+(1-k22)*g22);
%defining the probabilities when all are cooperators in the respective O_ks
cp = k1*k2*k11*k12*k21*k22;
c1 = P*k11*k12;
c2 = P*k21*k22;
c11 = P*k1;
c12 = P*k1;
c21 = P*k2;
c22 = P*k2;
%defining the probabilities when there is a defector in the respective U_ks
dU1 = (1-P);
dU2 = (1-P);
dU11 = (1-P)*(1-k1);
dU12 = (1-P)*(1-k1);
dU21 = (1-P)*(1-k2);
dU22 = (1-P)*(1-k2);

%replicator equations
dPdt = P*(1-P)*(cp*gop+gp-gop-xp+rhof);
```

```
    dk1dt = k1*(1-k1)*(c1*go1+(1-dU1)*(g1-go1-x1+rhof));
    dk2dt = k2*(1-k2)*(c2*go2+(1-dU2)*(g2-go2-x2+rhof));
    dk11dt = k11*(1-k11)*c11*(g11-x11+rhof);
    dk12dt = k12*(1-k12)*c12*(g12-x12+rhof);
    dk21dt = k21*(1-k21)*c21*(g21-x21+rhof);
    dk22dt = k22*(1-k22)*c22*(g22-x22+rhof);

    dzdt = [dPdt; dk1dt; dk2dt; dk11dt; dk12dt; dk21dt; dk22dt];
end
```
